# Exposure of gut bacterial isolates to the anthelminthic drugs, ivermectin and moxidectin, leads to antibiotic-like phenotypes of growth inhibition and adaptation

**DOI:** 10.1101/2024.01.17.575993

**Authors:** Julian Dommann, Jennifer Keiser, Julian Garneau, Alison Gandelin, Carlo Casanova, Peter M. Keller, Somphou Sayasone, Pascale Vonaesch, Pierre H. H. Schneeberger

## Abstract

Due to their broad-spectrum activities, ivermectin and moxidectin are widely used anthelminthics in veterinary and human medicine. However, ivermectin has recently been shown to perturbate gut-microbial growth. Given the macrolide-like structure of both ivermectin and moxidectin, there is a need to characterize the antibiotic spectrum of these anthelminthic drugs and their potential implications in the development of cross-resistance to macrolides and other families of antibiotics. Here, we incubated 59 bacterial isolates representing different clades frequently found in the gut with ivermectin and moxidectin at different concentrations for 16-72h. Further, we challenged 10 bacterial isolates with repeated and gradually increasing concentrations of these two anthelminthics and subsequently characterized their sensitivity to different antibiotics as well as ascending anthelminthic concentrations. We found, that antibacterial activity of the two anthelminthics is comparable to a selection of tested antibiotics, as observed by potency and dose dependence. Bacterial anthelminthic challenging *in vitro* resulted in decreased anthelminthic sensitivity. Further, adaptation to anthelminthics is associated with decreased antibiotic sensitivity towards three macrolides, a lincosamide, a fluoroquinolone, a tetracycline and two carbapenems. The observed change in bacterial sensitivity profiles is associated with - and likely caused by - repeated anthelminthic exposure. Hence, current and future large-scale administration of ivermectin and moxidectin, respectively, for the control of helminths and malaria raises serious concerns - and hence potential off-target effects should be carefully monitored.

## Background

Soil-transmitted helminths (STHs) are a multi-species group of parasitic, eukaryotic worms residing in species-specific niches within the host’s gastrointestinal (GI) tract. Amongst all neglected tropical diseases listed by the World Health Organization (WHO), STH infections dominate in terms of prevalence (over 1.5 billion) and disease burden [1, 2], mainly affecting pre-school and school-aged children and potentially leading to serious long-term health conditions, such as impairment in cognitive and physical development. While the improvement of water quality, sanitation and hygiene represents a long-term countermeasure, the short-term remedy relies on anthelminthic drugs to reduce morbidity [3–5]. The current frontline treatment encompasses the two benzimidazoles mebendazole and albendazole (ALB) [6]. However, cure rates vary in a species-specific manner [7, 8], with single-dose regimens of both drugs yielding poor cure rates in *Trichuris trichiura* infections [7, 9, 10]. As the pipeline for new drugs remains scarcely populated, the focus currently lies on repurposed veterinary drugs [11] and combination therapies [7, 8].

Ivermectin (IV) and moxidectin (MX) are two prime examples of such repurposed drugs. A 16-membered macrocyclic lactone ring, the chemical structure that represents the group of macrolide antibiotics, characterizes both IV and MX. Macrolide antibiotics such as erythromycin (EM), clarithromycin (CH) or azithromycin (AZ) interfere with bacterial protein synthesis [12] and are typically used in treatment of gram-positive bacterial infections. In contrast, IV and MX possess a broad activity spectrum against numerous parasites and are thus frequently administered as antiparasitic agents. MX was recently approved to treat human onchocerciasis [13] and was shown to have a good efficacy against *Strongyloides stercoralis* [14, 15] - and in combination with ALB - against *T. trichiura* [16]. IV on the other hand, is a widely used anthelminthic for filarial infections and used in combination therapy paired with ALB to increase treatment efficacy against *T. trichiura* infections [17], an indication recently added the WHO’s Model List of Essential Medicine [18].

As orally administered drugs pass through the small and large intestine, they are subject to interactions with numerous microbes before being absorbed, making the gut microbiome a relevant site for first-pass metabolism [19]. The pan-genome of gut microbes encompasses 150-fold more genes compared to their host, resulting in outstanding genetic diversity and therefore considerable possibilities of direct and indirect interactions that may influence bioavailability and efficacy of an administered compound [20].

*Vice versa*, drugs or their metabolites could also promote or inhibit bacterial growth, since antibacterial properties are not limited to antibiotic drugs [21]. Hence, orally administered non-antibiotic drugs could perturbate gut-microbial growth [21] and thus disrupt gut microbiome homeostasis. Due to the molecular structure of IV and MX, we hypothesize that various gut bacterial isolates could be sensitive to these drugs. Indeed, a recent study found an association between pre-treatment gut microbial composition and treatment outcome in participants receiving an IV-ALB combination therapy against *T. trichiura* infections [22]. Treatment failure was associated with gut bacterial species, such as *Prevotella copri, Streptococcus salivarius* and *Faecalibacterium prausnitzii* – among others [22]. Moreover, repeated IV or MX exposure could influence bacterial sensitivity profiles, which potentially leads to cross-resistant gut bacterial isolates or interferes with IV/MX treatment efficacy – depending on the underlying adaptation mechanism.

To our knowledge, only limited data regarding antibacterial activity of IV and MX against gut bacterial isolates have been published to date [21, 23]. A causal relationship between bacterial sensitivity profiles and IV or MX treatment would raise serious concerns in light of the extensive use of IV and MX in the livestock industry and the increase in distribution and administration for helminth infections and malaria transmission control [24–26]. Thus, it is crucial to anticipate potential off-target effects of these two anthelminthics. We therefore aimed to characterize antibacterial properties of these two widely used anthelminthics in-depth against a broad range of gut bacterial isolates *in vitro*, especially in the context of cross-reaction with other antibiotics.

## Results

### IV/MX inhibit *in vitro* growth of a broad range of gut bacterial isolates

To explore whether and to what extent IV/MX inhibit growth of gut bacteria, we conducted a series of *in vitro* experiments, incubating a broad taxonomic range of gut bacterial isolates in different concentrations of IV/MX. Specifically, we worked with 59 aerobically and anaerobically grown bacterial species (**Supplementary Tables 1-3**) across 10 genera (*Actinomyces, Bacteroides, Blautia, Clostridium, Dorea, Enterococcus, Escherichia, Lactobacillus, Staphylococcus, Streptococcus*). A detailed rationale covering the selection of isolates can be found in the **Methods** section. After oral administration, IV/MX immediately reach the target tissue for helminth infections – the gut linen –, resulting in estimated physiological concentration levels of approximately 5µM in the intestine [21]. We therefore incubated isolates with IV and MX at 1µM, 5µM and 10µM (**Supplementary** Figures 1 **– 10**). To compare growth curves across isolates and individual assays, we adapted the concept of area under the curve (AUC) ratios, as published by Gencay *et al.* [27]. Namely, we divided the AUC in presence of IV or MX at a given concentration, by the AUC of the corresponding control curve.

Overall, a drug concentration of 1µM IV/MX does not appear to have a negative impact on growth of any of the bacteria tested (**Figures 1A and 1B**). Inter-species comparison amongst aerobically incubated isolates reveals that isolates of the genus *Enterococcus* do not display delayed growth in presence of IV and MX across all tested concentrations (**Figure 1A**). In contrast, isolates belonging to the genera *Actinomyces* and *Streptococcus* are sensitive to the presence of IV and MX at 5 and 10 µM. Anaerobically, the tested isolates amongst the genera *Bacteroides, Lactobacillus and Staphylococcus* are not affected by the presence of either IV or MX (**Figure 1B**). In the contrary, we observed decreased AUC ratios amongst the genera *Blautia*, *Clostridium*, *Dorea* and *Streptococcus. Streptococcus salivarius (01)* and *Streptococcus parasanguinis (01)* were incubated both aerobically and anaerobically and were sensitive to IV/MX in both cases. Intra-species comparison amongst aerobically incubated isolates reveals drug dependent sensitivity. For instance, *S. pneumoniae (02)* displays lower sensitivity to IV or MX (AUC_IV10µM_: 0.84; AUC_MX10µM_: 0.47), compared *S. pneumoniae (01)* and *S. pneumoniae (03)* (AUC_IV10µM_: 0.47; AUC_MX10µM_: 0.1; AUC_IV10µM_: 0.1; AUC_MX10µM_: 0.1). Anaerobically, we also observed differences in intra-species sensitivity. In fact, 1/3 *Dorea* isolates (*D. longicatena (01);* AUC_IV5µM_: < 0.1) and 1/5 *Clostridium* isolates (*C. baratii (02);* AUC_MX10µM_: < 0.1) were inhibited by either IV or MX.

**Figure 1.**
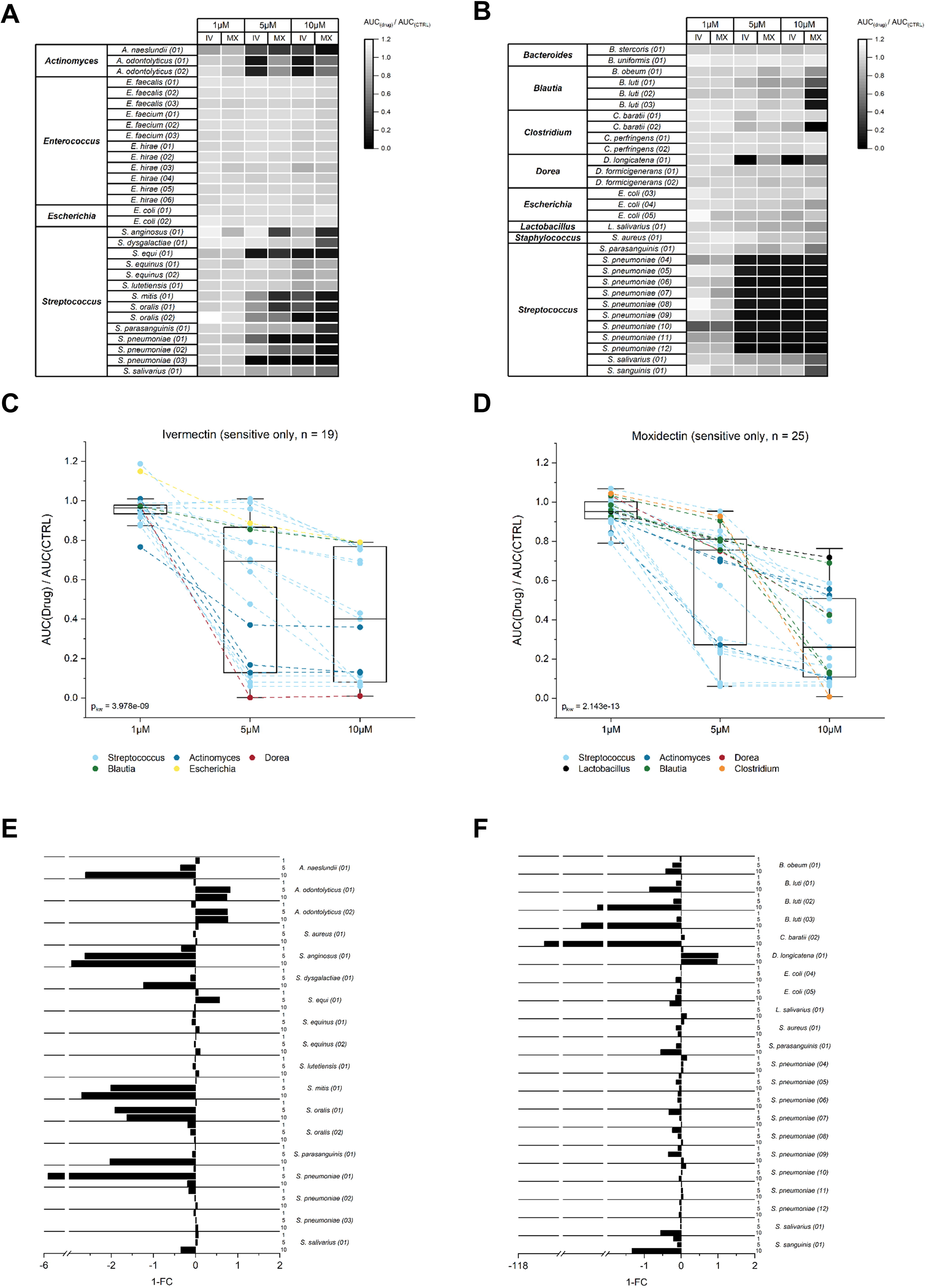
Sensitivity of gut bacterial isolates against ivermectin (IV) and moxidectin (MX): (A) AUC ratios of aerobically incubated isolates for IV and MX at concentrations of 1µM, 5µM and 10µM. (B) AUC ratios of anaerobically incubated isolates for IV and MX at concentrations of 1µM, 5µM and 10µM. (C) AUC ratios of 25 MX-sensitive isolates visualized across three concentrations (1µM, 5µM, 10µM). Values belonging to the same isolate are connected with a dotted line. Coloring was chosen according to the isolate genus. The boxplot whiskers indicate the 95% CI. (D) AUC ratios of 19 IV-sensitive isolates visualized across three concentrations (1µM, 5µM, 10µM). Values belonging to the same isolate are connected with a dotted line. Coloring was chosen according to the isolate genus. The boxplot whiskers indicate the 95% CI. (E, F) Fold change (FC) ratio for sensitive isolates. FC was calculated as 1 - (AUC_IV_/AUC_MX_).l – FC > 0 indicates a higher sensitivity towards IV, 1 – FC < 0 indicates higher sensitivity towards MX. The figure is stratified by aerobic (E) and anaerobic (F) incubations.

To transition from quantitative values (AUC ratio) to qualitative values in Figure 1C and 1D (“sensitive”; “not sensitive”) we employed a tentative AUC ratio cut-off of 0.8 to stratify isolates into “sensitive” and “not sensitive”. Pairwise comparison of isolates across the three concentrations resulted in an inverse relationship between drug dose and AUC ratio (dose dependence; **Tables 1-2**).

According to the tentative cut-off, 25/59 isolates were considered sensitive towards MX and 19/59 towards IV. Although there is a general trend of lower AUC ratios with higher drug concentrations for both IV and MX, AUC ratios vary substantially across genera and species in presence of 5µM or 10µM IV or MX. Lastly, we also observed a phenotype consisting of a drug-specific response in several isolates. For instance, incubation of *S. anginosus (01)* and *S. dysgalactiae (01)* (**Figure 1A and 1E**) with MX results in lower AUC ratios, compared with IV at the same concentration. In contrast *A. odontolyticus (01), A. odontolyticus (02)* and *D. longicatena (01)* seem to have a higher sensitivity towards IV. Within our set of tested species, more isolates appear to be sensitive towards MX (**Figure 1E and 1F**) and at equal concentrations of IV or MX, MX results in lower AUC ratios (**Figure 1A and 1B**).

### *In vitro* potency of IV/MX is comparable to a selection of antibiotics

We next sought to compare the potency of the two anthelmintic drugs to that of antibiotic compounds *in vitro.* Since IV and MX belong to the macrocyclic lactone drug class, we included the three macrolide antibiotics EM, CH and AZ in the comparison. Clindamycin (CL) represents a lincosamide antibiotic, which shares the mode of action (MOA) with macrolide antibiotics. We further included the fluoroquinolone ciprofloxacin (CX), tetracycline (TC) and the two carbapenems, meropenem (MP) and imipenem (IP), as they possess completely different modes of action. Therefore, we incubated a subset of anthelmintic-susceptible bacterial isolates, comprised of 8 *Streptococcus spp.* and 3 *Actinomyces spp.* identified in our primary screen, with a selection of antibiotics (EM, CH, AZ, CL, CX, TC, MP, IP) at concentrations of 1µM, 5µM and 10µM to simulate the magnitude as in incubations with IV/MX. In contrast to IV and MX, the tested antibiotics cause a reduction in AUC ratios at 1μM (**Supplementary** Figure 1, KW; p **=** 1.186e-08). However, incubations with IV/MX at 5µM or 10µM result in AUC ratios in a comparable range as the tested antibiotics (KW; p = 0.03708 and KW; p = 0.8284). *S. dysgalactiae (01)* represents an exception at 5µM, as it displays high AUC ratios for both IV and MX (**Supplementary** Figure 2, AUC_IV5µM_: 1.06; AUC_MX5µM_: 0.95). Similarly, solely *S. dysgalactiae (01)* displays a high AUC ratio for IV at a concentration of 10µM (**Figure 2**, AUC_IV10µM_: 0.96). In the case of *S. oralis (02)* and *S. pneumoniae (03)*, IV and MX at 10µM achieved the lowest AUC ratios across all tested drugs (AUC_IV10µM_: 0.07; AUC_MX10µM_: 0.07 and AUC_IV10µM_: 0.06; AUC_MX10µM_: 0.06). Results of a pairwise Wilcoxon rank sum test between all compounds can be found in **Supplementary Tables 5-7**. Furthermore, we performed a Pearson correlation (**Supplementary Tables 8-13**) based on AUC ratios of the 11 isolates in presence of all macrolide (IV, MX, EM, CH, AZ) and lincosamide (CL) compounds for each tested concentration. There, we could observe pairwise correlation within macrolide antibiotics (EM, CH, AZ) for all tested concentrations, but not between anthelminthics (IV, MX) and macrolide (EM, CH, AZ) or lincosamide (CL) antibiotics.

**Figure 2.**
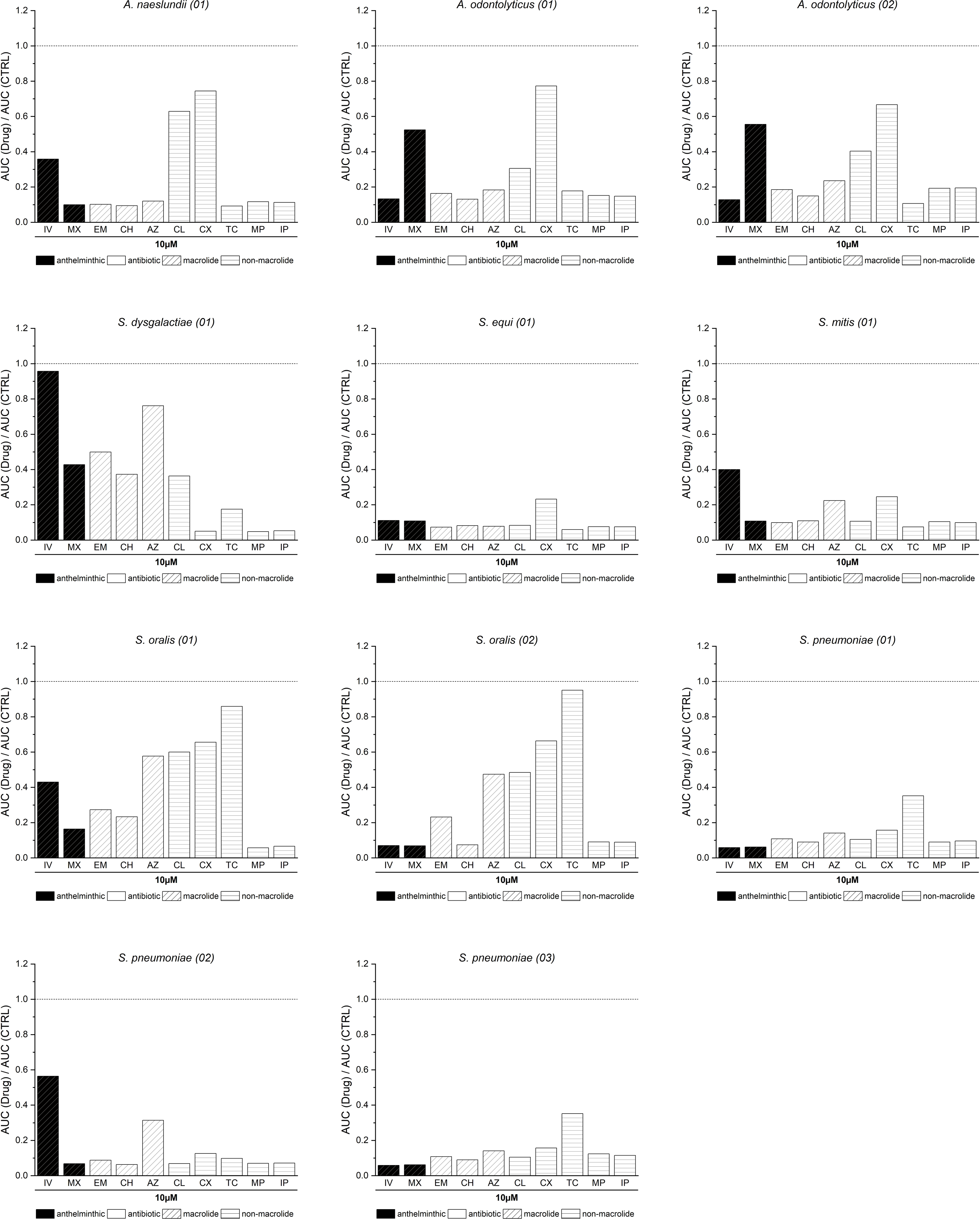
In vitro potency of ivermectin (IV) and moxidectin (MX): AUC ratios of 11 bacterial isolates in presence of IV, MX, EM, CH, AZ, CL, CX, TC, MP or IP at lOµM. The dotted line marks theoretical uninhibited growth of the isolate (AUC ratio = 1). Bar colors correspond to the primary usecase of the compound. Black = anthelminthic, white = antibiotic.

### Bacterial isolates display adaptation to IV/MX and a selection of antibiotics after anthelminthic challenging experiments

Since IV and MX inhibit bacterial growth in a broad range of isolates, we aimed to test whether sensitive bacterial isolates can adapt to repeated sub-inhibitory exposure of IV and MX and whether this adaptation changes the antibiotic susceptibility profiles of these isolates. Five bacterial isolates, namely *S. salivarius (01), S. parasanguinis (01), S. pneumoniae (02), S. mitis (01)* and *S. dysgalactiae (01)*, and their challenged counterparts (appendices “-IVc” or “-MXc”) were incubated with IV/MX and antibiotics as described before (**Supplementary** Figures 13-17). As the primary focus of these incubations was to solely demonstrate the effect of pre-challenging on antibiotic sensitivity, we opted for sub-lethal doses (i.e. 0.1µM or 0.01µM) if no growth could be measured at concentrations of ≥ 1µM.

Comparing the IV and MX AUC ratios of the original to the challenged isolates, we observed three phenotypes (**Figure 3**). First, we documented increased AUC ratios for both IV and MX in response to anthelminthic challenging with either drug. For instance, for *S. salivarius (01-IVc)* (AUC_IV10µM_: 0.73; AUC_MX10µM_: 0.60) compared to *S. salivarius (01)* (AUC_IV10µM_: 0.68; AUC_MX10µM_: 0.51). We also noted different AUC ratios in the presence of either anthelminthic at 10μM for *S. salivarius (01-MXc)* (AUC_IV10µM_: 0.77; AUC_MX10µM_: 0.91) compared to *S. salivarius (01)*. This phenotype is shared with *S. pneumoniae (02-IVc)* and *S. pneumoniae (02-MXc)*, *S. mitis (01-IVc)*, and *S. mitis (01-MXc)* and lastly *S. dysgalactiae (01-IVc)* compared to their respective unchallenged original isolate. In contrast we observed a different phenotype in *S. parasanguinis (I01)*, namely that AUC ratios only increase when re-challenged with the same anthelminthic. For instance, *S. parasanguinis (01-IVc)* (AUC_IV10µM_: 0.96; AUC_MX10µM_: 0.25) possesses a higher AUC ratio in presence of IV compared to *S. parasanguinis (01)* (AUC_IV10µM_: 0.79; AUC_MX10µM_: 0.26), but a lower AUC ratio in presence of MX. Similarly, *S. parasanguinis (01-MXc)* (AUC_IV10µM_: 0.75; AUC_MX10µM_: 0.74) displays higher AUC ratios compared to *S. parasanguinis (01)* in the presence of MX, compared to lower ratios in presence of IV. Lastly, *S. dysgalactiae (01-MXc)* (AUC_IV10µM_: 0.84; AUC_MX10µM_: 0.19) exhibits a third and unique phenotype displaying lower AUC ratios compared to *S. dysgalactiae (01)* for both IV and MX incubations.

**Figure 3.**
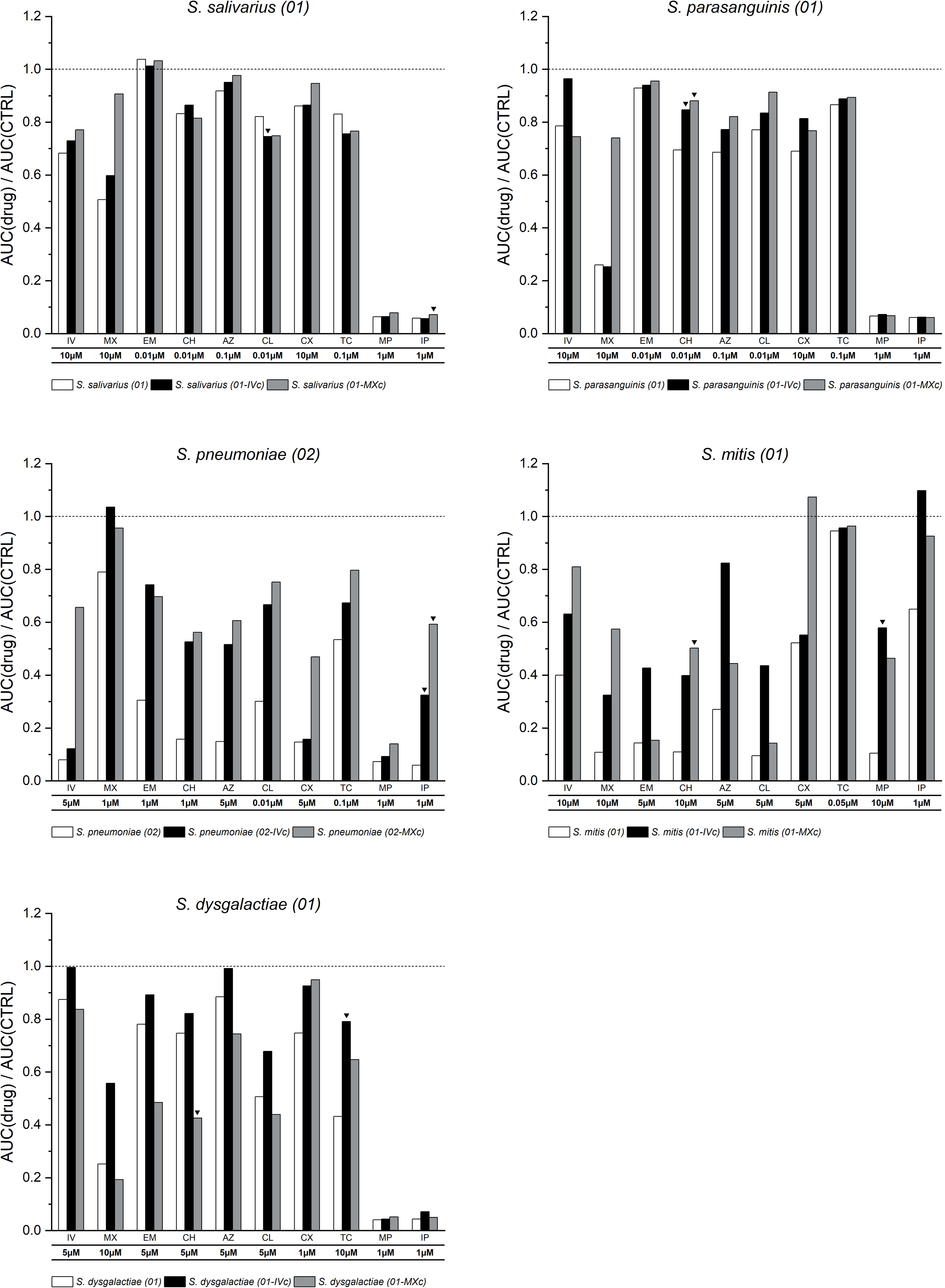
Adaptation of bacterial isolates in response to anthelminthic challenging: AUC ratios of 10 challenged bacterial isolates and their original counterparts in presence of IV, MX, EM, CH, AZ, CL, CX, TC, MP or IP at different concentrations. The dotted line marks theoretical uninhibited growth of the isolate (AUC ratio = 1). Bar colors correspond to the challenging status. White = original isolate, black = IV-challenged isolate, grey = MX-challenged isolate. For each challenged isolate, we highlighted the compound resulting in the highest \1-FC\ (black triangles).

To compare the effects of anthelminthic-challenging on antibiotic sensitivity, we used the fold-change (FC) of AUC ratios of the original isolate divided by the AUC ratio of the challenged isolate, for each compound individually. We considered that antibiotic sensitivity was affected when (1-FC) ≥ 0.1. Following exposure to MX, we observed a change in sensitivity towards MP and IP in *S. salivarius (01)* and the highest fold-change for IP (|1-FC_IP1µM_| = 0.20). These changes were not observed following exposure to IV, except for CL (|1-FC_CL0.01µM_| = 0.10). For *S. parasanguinis (01-IVc)*, we observed a change in sensitivity in presence of CH, AZ and CX. For *S. parasanguinis (01-MXc)* we observed an equal pattern with the addition of CL. We observed the highest fold-change for *S. parasanguinis (01-IVc)* and *S. parasanguinis (01-MXc)* in presence of CH (|1-FC_CH0.01µM_| = 0.18 and 0.21). Prolonged exposure to MX in our tested *S. pneumoniae* isolate was associated with a decrease in sensitivity to all tested antibiotics (range of (highest fold change: |1-FC_IP1µM_| = 0.89). We observed a similar result following exposure to IV, except for the antibiotic CX to which sensitivity of the challenged isolate was like that of the unchallenged isolate (highest fold change: |1-FC_IP1µM_| = 0.82). In the case of *S. mitis (01-IVc)*, sensitivity towards all antibiotics except CX and TC decreased, and this decrease was the highest for MP (|1-FC_MP1µM_| = 0.82) resulting in almost uninhibited growth of the challenged isolate. For *S. mitis (01-MXc),* antibiotic sensitivity was found to be the lowest for CH (|1-FC_CH10µM_| = 0.78) but did not change for EM and TC. For *S. dysgalactiae (01)*, some of the changes observed post-exposure were inverted, meaning that the unchallenged isolate was less sensitive to another antibiotic than the anthelmintic-challenged one. For instance, MX-challenging was associated with increased sensitivity to the three macrolide antibiotics (EM, CH, AZ) and CL (highest fold change: |1-FC_CH5µM_| = 0.75). This was not the case for the isolate that was challenged with IV (highest fold change: |1-FC_TC10µM_| = 0.45), suggesting different adaptation mechanisms in this isolate. Sensitivity was found to be higher against CX and TC but remained unchanged for MP and IP, irrespective of the anthelmintic drug used for challenging.

## Discussion

A handful of studies have described some antibacterial properties of IV [28–30]. In this study, we systematically tested antibacterial properties of IV - and the closely related compound MX - against 59 (gut) bacterial isolates spanning across 10 genera and originating from both clinical and commercial sources. We included lincosamide and macrolide resistant clinical isolates (20/59) to test whether macrolide or lincosamide resistance is coupled to sensitivity to IV/MX. We further included clinical isolates (27/59) enriched from stool samples obtained in a study in Lao PDR, due to their extensive clinical background. Lastly, we included several commercial strains (12/59) to broaden the taxonomic range of our results. We found that both IV and MX inhibit growth of a broad range of bacterial isolates *in vitro* in a dose dependent manner with comparable potency. Macrolide and lincosamide antibiotics achieve their bacteriostatic or bactericidal activity against gram-positives by binding to the 50S subunit of the bacterial ribosome. For IV and MX no MOA against bacteria has been identified yet. Given our data however, we can speculate that the MOA of IV and MX seems to be different from macrolide or lincosamide antibiotics, as, first, we observed different sensitivity profiles of bacterial isolates in response to the macrolides EM, CH and AZ and the lincosamide CL compared to IV and MX (**Supplementary Tables 8-13**). Secondly, we observed changes in the bacterial sensitivity profile of the macrolide or lincosamide resistant isolates *S. pneumoniae (02)*, *S. dysgalactiae (01)* and *S. mitis (01)* after anthelminthic challenging, indicating a further adaptation (different MOA), opposed to a redundant one (same MOA).

We demonstrated adaptation to IV and MX, associated with decreased sensitivity *in vitro*. A possible shortcoming of our study is that aggressive *in vitro* challenging might not be comparable to real-world applications of IV and MX. To control STH infections, MDA is usually conducted once to twice a year. According to Maier *et al.*, 8mg of MX and 200µg/kg IV (i.e., the standard dose used for helminth infections) result in estimated intestinal concentrations of 4.2 or 4.6μM, respectively [21]. However, IV was recently considered as a vector control drug against female *Anopheles spp.* mosquitos, potentially introducing more frequent and higher doses [25, 31]. Furthermore, the recent MORDOR trial demonstrated increased macrolide resistance within the gut microbiome already after administration of the macrolide antibiotic AZ twice a year for 2 years [32, 33].

We observed broad adaptation phenotypes for *S. salivarius (01)*, *S. pneumoniae (02)* and *S. mitis (01)*, as challenging with anthelminthics resulted in decreased sensitivity against both anthelminthics. In contrast, anthelminthic challenging of *S. parasanguinis (01)* resulted in decreased sensitivity only against the corresponding anthelminthic, suggesting a drug-specific mechanism, opposed to a potentially broad mechanism as adopted by *S. salivarius (01)*, *S. pneumoniae (02)* and *S. mitis (01).* Interestingly, in the macrolide and lincosamide resistant isolate *S. pneumoniae (02)*, anthelminthic challenging seemed to exert a broad-spectrum effect as both, IV- and MX-challenged isolates displayed decreased sensitivity towards all antibiotic classes it was tested against (macrolides, lincosamide, tetracycline, fluoroquinolone, and carbapenem). In addition, this broad-spectrum interaction between anthelminthic and macrolide or lincosamide resistance was also observed in *S. mitis (01-IVc)*, *S. mitis (01-MXc)* and *S. dysgalactiae (01-IVc).* Hence, further characterization of resistance mechanisms is crucial to understand the development and maintenance of AMR in clinically relevant species. This is specifically true for *S. pneumoniae* which is one of the six leading pathogens causing lethal lower respiratory infections [34]. *S. pneumoniae* is primarily found in the upper respiratory tract, therefore, in its primary niche, contact with high concentrations of IV or MX is unlikely. However, dynamic transfer of resistance genes across the microbiome was recently demonstrated [35] and would a possible scenario in the case of IV/MX MDA. It is also critical to investigate this interaction in other commensal species as they might act as reservoir for AMR genes and thus contribute to the persistence of AMR in the human body. Populations at risk for helminth infections and other infectious diseases often overlap in the Global South. For instance, sub-Saharan Africa currently is attributable to most deaths related to AMR [34]. There, anthelminthics could further contribute to the AMR in bacteria. Additionally, both Southeast Asia and India represent hotspots of helminth prevalence [36] and antibiotic use [37] – providing a potential environment for the described dynamics between antibiotic and resistant bacteria. Therefore, this broad-spectrum effect anthelminthic challenging in antibiotic resistant bacteria is not to be neglected and should be investigated in further studies.

Our study presents few limitations. Firstly, it does not uncover the mode of action of IV and MX in bacteria and corresponding adaptation mechanisms on a molecular level. In further studies, we aim to conduct genomic and transcriptomic analyses to fill this gap. Moreover, our experiments did not test for long-term adaptation. Thus, the described adaptation could be transient. This shortcoming should also be addressed in further experiments.

In conclusion, our study yielded three key novel findings: i) exposure to IV and MX causes phenotypical adaptation, ii) adaptation mechanisms are likely diverse, encompassing both drug-specific and broad-spectrum mechanisms, and iii) adaptation to IV/MX also imparts varying degrees of decreased sensitivity to other antibiotics. Both IV and MX remain crucial compounds to combat several infectious diseases. However, in the context of the uprising of IV and MX, this raises serious concerns, as repeated anthelminthic exposure of gut-bacterial isolates may trigger broad cross-resistant phenotypes, potentially interfering with vital antibiotic treatments.

## Methods

### Origin of bacterial isolates

20/59 bacterial isolates are macrolide or lincosamide resistant and were obtained via the Institute for Infectious Diseases (Berne, Switzerland). As lincosamides share the mode of action with macrolides, resistance mechanisms often overlap [38]. We therefore included these isolates to test whether macrolide or lincosamide resistance is further coupled to sensitivity to IV/MX. The lincosamide resistant strains were comprised of 5 *Streptococcus spp.* and *3 Actinomyces spp. Streptococcus* species are frequent, gram-positive commensals in the oral microbiota, but are especially abundant in the small intestine [39]. However, several mechanisms of macrolide resistance have been identified among *Streptococcus* species, including mainly drug efflux, target alteration, and drug inactivation [40–42]. Moreover, *Streptococcus salivarius* – amongst others – was associated with ALB-IV combination treatment failure in a recent study [22]. *Actinomyces* spp. on the other hand are close relatives to avermectin producing bacteria (*Streptomyces avermitilis*). They therefore likely possessed unique co-incubation phenotypes with derivatives such as IV or MX. The macrolide resistant isolates were comprised solely of *S. pneumoniae* isolates (n = 12). *S. pneumoniae* is primarily found in the lungs but also commonly found throughout the upper and lower GI tract, and is one of the six leading pathogens causing lethal lower respiratory infections [34]. We included 27/59 bacterial isolates from clinical stool samples from a recent study conducted in Lao PDR (NCT03527732). As Lao PDR represents a prime site of clinical trials involving both IV and MX and lies in midst of a hotspot of helminth prevalence, the corresponding bacterial isolates likely reflect an extensive clinical background. Bacterial isolation from stool samples was either conducted at Swiss TPH in Allschwil, Switzerland (18/59 bacterial isolates) or by Dr. Julian Garneau and Alison Gandelin at the University of Lausanne in Lausanne, Switzerland (9/59 bacterial isolates). Enriched clinical isolates were comprised of the genera *Blautia, Clostridium, Dorea, Enterococcus, Escherichia* and *Streptococcus.* It is critical to investigate this interaction with IV/MX in several commensal species as they might act as reservoir for AMR genes and thus contribute to the persistence of AMR in the human body. Therefore, we included 12/59 commercial isolates (*Bacteroides, Blautia, Dorea, Lactobacillus, Staphylococcus, Streptococcus*), to further broaden taxonomic range of our results. The commercial isolates were purchased from the German Collection of Microorganisms and Cell Cultures (https://www.dsmz.de/). We additionally generated 10 anthelminthic challenged isolates through a procedure described later. We opted for *Streptococcus* species due to reasons mentioned above. Moreover, we included lincosamide resistant (*S. mitis, S. dysgalactiae*), macrolide resistant (*S. pneumoniae*) and commercial strains (*S. salivarius, S. parasanguinis*) to characterize the phenotype of anthelminthic challenging in these differing genotypes. An overview of the challenged bacterial isolates is given in **Supplementary Table 4**.

### Enrichment and identification of clinical isolates

Clinical stool samples were obtained from a recent study in Lao PDR (NCT03527732) [40]. For each sample, 50-100mg of stool were homogenized in 500μl BHI + 5% yeast in a safety cabinet (aerobic conditions). After centrifugation (1min, 15000rcf), 80µl of the supernatant was added to 10ml of the corresponding enrichment medium (**see Supplementary Table 2**) and left overnight in a 5% CO_2_ incubator at 37°C. The following culture media were used: Brain heart infusion broth (BHI; Thermo Fisher Scientific, CM1135), yeast extract (Thermo Fisher Scientific, LP0021B), inulin (Thermo Fisher Scientific, 457100250), dehydrated Todd-Hewitt broth (TH; Thermo Fisher Scientific, CM0189), modified gifu anaerobe medium (mGAM; HyServe, 05433), dehydrated Schaedler broth (Thermo Fisher Scientific, CM0497B) and Lennox broth (LB; Thermo Fisher Scientific, 12780052). The next day, each culture was diluted 1:1000 in BHI + 5% yeast and 10μl were streaked on a BHI + 5% yeast agar (Thermo Fisher Scientific, CM1136) plate. The streaked agar plates were left overnight in a 5% CO_2_ incubator at 37°C. The following day, individual colonies were randomly picked and re-inoculated in fresh BHI + 5% yeast. Again, cultures were left to grow overnight in a 5% CO_2_ incubator at 37°C. Finally, 20% glycerol stocks were prepared of each isolate and stored at - 80°C. To identify the isolated bacteria, Columbia agar plates with 5% sheep blood (Thermo Fisher Scientific, PB5039A) were streaked with the corresponding glycerol stocks. Colonies were left to grow overnight in a 5% CO_2_ incubator at 37°C. The next day, MALDI-TOF analysis was performed by the Institute of Medical Microbiology (University of Zürich, Switzerland) to identify the isolates. In brief, single colonies are loaded onto a MALDI-TOF steel target plate using a sterile toothpick. Each colony on the plate is treated with 1μl of 25% formic acid and subsequently with 1μl of α-Cyano-4-hydroxycinnamic (CHAC) matrix. After sample preparation, the plate is loaded into a Microflex MALDI-TOF instrument for spectra measurement (Bruker Daltonics) and subsequent identification [43].

### Cultivation of bacterial isolates

Bacterial glycerol stocks were stored at -80°C. To cultivate an isolate, the glycerol stock was first thawed on ice. Subsequently, 10μl of the thawed glycerol stock was used to start a culture in 10ml of culture medium and left to grow at 37°C. BHI + 5% yeast was used exclusively to cultivate all bacterial isolates. The incubation period for all isolates was between 24h – 72h. Anaerobic cultivation was performed in a vinyl anaerobic chamber (Coy Laboratory Products, Michigan, United States) with a gas mix composed of 85% N_2_, 10% CO_2_ and 5% H_2_. Inoculated isolates were grown in the integrated anaerobic incubator at 37°C. Aerobic inoculations were performed in a safety cabinet (SKAN Berner, Elmshorn, Germany). Inoculated aerobic isolates were grown in a 5% CO_2_ incubator (Binder, Tuttlingen, Germany) at 37°C. To preserve bacterial cultures after cultivation, 100μl of culture was added to 100μl of 40% glycerol solution (Thermo Fisher Scientific, 17904) in purified water. The resulting 20% bacterial glycerol stocks were frozen and stored at -80°C until further use.

### Bacterial anthelminthic challenging experiments

Five bacterial isolates - originally *S. salivarius (01)*, *S. parasanguinis (01)*, *S. pneumoniae (02)*, *S. mitis (01)* and *S. dysgalactiae (01)* - were challenged with increasing concentrations of anthelminthics (IV, MX) up to a final concentration of 20μM, resulting in IV or MX challenged isolates *S. salivarius (01-IVc)*, *S. salivarius (01-MXc)*, *S. parasanguinis (01-IVc)*, *S. parasanguinis (01-MXc)*, *S. pneumoniae (02-IVc)*, *S. pneumoniae (02-MXc), S. mitis (01-IVc), S. mitis (01-MXc), S. dysgalactiae (01-IVc) and S. dysgalactiae (01-MXc)* (**Supplementary Table 4**). In brief, each isolate was initially cultured in liquid growth media (BHI + 5% Yeast) supplemented with 5μM IV or MX for 24h. After 24h, 100µl of the grown culture was passaged to fresh growth medium supplemented with 2-fold higher anthelminthic concentration (i.e. 10µM) and incubated again for 24h. This cycle was continued repeatedly until a final challenging concentration of 20μM is reached (3 cycles). If an isolate failed to grow, it was left for another 24h (total incubation time = 48h). This was the case for *S. dysgalactiae (01)* in 20μM MX and acti_odo_02 in 20μM IV. For *S. pneumoniae (02)* and *S. mitis (01)* the only a final MX concentration of 10μM was reachable (2 cycles). 20% glycerol stocks were prepared for all isolates and each round of a cycle. The glycerol stocks were stored at -80°C until further use. To confirm identity of the isolate and exclude contamination, potentially introduced by long incubation periods, the final challenged isolates underwent 16S rRNA gene sequencing on the ONT MinION platform using the ONT Native Barcoding Kit (SQK-NBD114.24) on a Flongle Flow Cell (FLO-FLG114). The resulting sequences were identified using Emu [44] equipped with its full-length 16S database.

### *In vitro* incubation assays with anthelminthic or antibiotic drugs

Throughout this study, bacterial isolates were incubated with a variety of anthelminthics (IV, MX) or antibiotics, namely erythromycin (EM; Lubio, ORB322468), clarithromycin (CH; Lubio, HY-17508), azithromycin (AZ; Lubio, ORB322360), clindamycin (CL; Lubio, HY-B0408A), tetracycline (TC; Lubio, HY-B0474), ciprofloxacin (CX; Lubio, HY-B0356A), meropenem (MP; Lubio, A5124) and imipenem (IP; Lubio, ORB1308410). Incubations were carried out in a 96-well plate format to facilitate optical density measurements (at λ = 600nm; OD600) using a Hidex Sense plate reader (Hidex Oy, Turku, Finland). OD600 was measured every 30-60min at 37°C for 16-72h. Drug solutions were prepared as follows: First, the corresponding drug amounts were weighed ± 0.1mg on an Ohaus Adventurer AX124 analytical scale (Ohaus, Nänikon, Switzerland) and dissolved in the according volume of solvent to generate a 5000μM working solution (WS). We used UltraPure™ DNase/RNase-free distilled water (Thermo Fisher Scientific, 10977-035) to dissolve CL, CX, TC and IP. Dimethylsulfoxide (DMSO; Sigma-Aldrich, 41640) was used to dissolve IV, MX, EM, CH, AZ and MP. 2500μM and 500μM WS were obtained by diluting the 5000μM WS. Working solutions were prepared fresh each week and kept at -20°C while limiting the amount of freeze and thaw cycles. WS only represent an intermediate step and had to be diluted in BHI + 5% yeast to 1.25x of the desired final assay concentration (12.5μM, 6.25μM, 1.25μM). Prior to the incubation, the bacterial isolates were cultivated as described before. In the case of challenged isolates, 20μM of either IV or MX - or 10μM in the case of *S. pneumoniae (02-MXc)* and *S. mitis (01-MXc)* - was added, to replicate the former cultivation environment. Grown bacterial cultures were diluted 1:200 in BHI + 5% yeast, before combining them with the corresponding 1.25x drug solutions. In the case of challenged isolate cultures, the dilution was 1:20’000. Each well of a plate contained a ratio of bacterial culture : 1.25x drug solution of 1 : 4. The final 1x drug concentration was therefore reached and DMSO concentration was constant at 0.2%. During each incubation, we added two control per isolate: As some drugs needed to be dissolved in DMSO – which is known to inhibit bacterial growth in higher concentrations [45] - a DMSO control (isolate in 0.2% DMSO) was added to each plate. A second control represented a positive control (PC), meaning the isolate was added to growth medium only. We incubated 28/59 isolates under anaerobic conditions as they either did not tolerate oxygen (strict anaerobes) or were enriched under anaerobic conditions. Likewise, 29/59 isolates were incubated under aerobic conditions. The remaining 2/59 isolates (*S. salivarius, S. parasanguinis*) we incubated in both conditions, to test whether the resulting phenotypes would differ.

### Bacterial DNA extraction

To extract bacterial DNA from liquid cultures, we used the DNeasy PowerSoil Pro kit (QIAGEN, 47016) and followed the manufacturer’s standard protocol included in the kit using 100μl of the bacterial culture. Final DNA concentrations were determined using a QuBit^TM^ 4 fluorometer (Thermo Fisher Scientific, Q33238) with the 1x dsDNA HS Assay kit (Thermo Fisher Scientific, Q33231). Eluates were stored at -20°C until further use.

### Library preparation for bacterial 16S rRNA sequencing

Bacterial 16S rRNA sequencing was performed on the MinION Mk1C platform (Oxford Nanopore Technologies, United Kingdom of Great Britain). All samples were processed according to the manufacturer’s instructions provided with the 1-24 16S barcoding kit (Oxford Nanopore Technologies, SQK-16S024). In brief, the DNA concentration of all samples was measured using a QuBit^TM^ 4 fluorometer (Thermo Fisher Scientific, Q33238) and the 1x dsDNA HS Assay kit (Thermo Fisher Scientific, Q33231). Samples were diluted using UltraPure™ DNase/RNase-free distilled water (Thermo Fisher Scientific, 10977-035) to reach a final input of 25ng DNA. Next, the samples were barcoded and amplified on a CFX Opus 384 Real-Time PCR system (Biorad, Cressier, Switzerland). Subsequently, the barcoded amplicons were cleaned up in multiple steps using the AMPure XP reagent (Beckman Coulter, A63881) in combination with the DynaMag^TM^-2 magnetic tube stand (Thermo Fisher Scientific, 12321D) and a HulaMixer^TM^ (Thermo Fisher Scientific, 15920D). Finally, the sequencing libraries were pooled and sequenced using a flongle flow cell (Oxford Nanopore Technologies, FLG-001). Sequencing was performed for approximately 24h.

### Agarose gel electrophoresis

After the amplification step within the 1-24 16S barcoding kit protocol, gel electrophoresis was performed to visualize successful 16S rRNA gene amplification. A 1% agarose solution in 1x Tris-acetate-EDTA (Thermo Fisher Scientific, B49) was prepared using analytical grade agarose powder (Promega, V3121). The GeneRuler 1kb Plus DNA Ladder (Thermo Fisher Scientific, SM1331) and samples were combined with 6x DNA Loading Dye (Thermo Fisher Scientific, R0611) in a 5:1 ratio (sample:dye). Gel electrophoresis was running in 1x TAE for 25min at 100V. For final staining, the gel was bathed in a GelRed® nucleic acid gel stain solution (Biotium, 41003) for 20min. UV imaging was performed using a Lourmat Quantum ST5 Imager (Vilber, Collégien, France). Images were taken using the VisionCapt software (v15.0) with an exposure time of 2-3s.

### Data Analysis

Data of bacterial growth curves, as well as area under the curve (AUC) ratios were curated in Microsoft Excel 2016 and plotted using OriginPro 2022b (OriginLab Corporation, Northhampton MA, United States). To calculate AUC ratios the R package growthcurver [42] (v0.3.1) was used in RStudio equipped with R 4.1.3. All original growth curves that were used for AUC ratio calculation can be found in the Supplementary Material.

### Statistics

To test for dose dependence in IV/MX incubations, we applied a pairwise Wilcoxon rank sum exact test (to test AUC ratio pairs for two concentrations; p-value adjustment method = BH) and a kruskal wallis test (to test corresponding AUC ratios across three concentrations; degrees of freedom = 2) in RStudio equipped with R 4.1.3 to the AUC ratio values of IV/MX-sensitive isolates (**Tables 1-2**). To test, for each incubation concentration, whether potency of the antheminthics is comparable to antibiotics, we utilized a kruskal wallis test (degrees of freedom = 1) with the data of 11 bacterial isolates (**Figure 2, Supplementary** Figures 1-2). P-values for pairwise comparisons of anthelminthic and antibiotic compounds (**Supplementary Tables 5-7**) were obtained via a Wilcoxon rank sum exact test (p-value adjustment method = BH). The Pearson correlation matrix and p-values for AUC ratios of 11 bacterial isolates in presence of IV/MX or macrolides/lincosamide antibiotics were calculated in Microsoft Excel 2016.

**Table 1.** Dose dependence in IV incubations: P-values of pairwaise Wicoxon rank sum tests (W) and kruskal-wallis (KW) test performed on AUC ratios of IV-sensitive bacterial isolates for the incubation concentrations 1µM, 5µM and 10µM (IV = ivermectin).

|  | IV |  |  |  |
| --- | --- | --- | --- | --- |
| Concentrations | 1µM vs. 5µM (W) | 5µM vs.10µM (W) | 1µM vs.10µM (W) | 1µM vs.5µM vs.10µM (KW) |
| p-value | 5.40E-05 | 1.30E-01 | 3.20E-09 | 4.08E-07 |

**Table 2.** Dose dependence in MX incubations: P-values of pairwaise Wicoxon rank sum tests (W) and kruskal-wallis (KW) test performed on AUC ratios of MX-sensitive bacterial isolates for the incubation concentrations 1µM, 5µM and 10µM (MX = moxidectin).

|  | MX |  |  |  |
| --- | --- | --- | --- | --- |
| Concentrations | 1µM vs. 5µM (W) | 5µM vs.10µM (W) | 1µM vs.10µM (W) | 1µM vs.5µM vs.10µM (KW) |
| p-value | 2.10E-08 | 6.00E-04 | 4.70E-14 | 2.86E-11 |

## Acknowledgements

We thank the Lao TPHI team, local authority, field research team and study participants for facilitating the clinical trial NCT03527732.We are thankful to Diana Albertos Torres (Institute of Medical Microbiology, University of Zürich, Switzerland), who kindly agreed to identify 18 bacterial colonies isolated from stool by MALDI-TOF-MS. We are grateful to the European Research Council (No. 101019223) for financial support.

## Data Availability

The sequencing data (16S rRNA gene sequencing) generated in this study have been deposited in the NCBI Short Read Archive under the accession PRJNA1053597.

## Author contributions

J.D.: study design, research design, project supervision, experimental work, statistical analyses, figure generation, writing of the initial paper, and paper editing. J.K.: study design, research design, project supervision, funding acquisition and paper editing. S.S: study design, research design, project supervision, field implementation. J.G. and A.G.: experimental work (enrichment of clinical isolates), paper editing. C.C., P.M.K. and P.V.: paper editing. P.H.H.S.: study design, research design, project supervision, and paper editing.

