## Supplementary Material (PDF) for "Exposure of gut bacterial isolates to the anthelminthic drugs, ivermectin and moxidectin, leads to antibiotic-like phenotypes of growth inhibition and adaptation"

*Supplementary Table 1: Overview of the 20 macrolide (erythromycin) or lincosamide (clindamycin) resistant bacterial isolates used in the study.*

| isolate | sample type | resistance (MIC[mg/l]) |
|---|---|---|
| *Actinomyces naeslundii (01)* | blood cluture | clindamycin (>256) |
| *Actinomyces odontolyticus (01)* | wound swab (cervical) | clindamycin (>256) |
| *Actinomyces odontolyticus (02)* | peritonsillar abscess swab | clindamycin (>256) |
| *Streptococcus dysgalactiae (01)* | synovial fluid knee | clindamycin (>256) |
| *Streptococcus equi (01)* | periprosthetic tissue biopsy | clindamycin (1.5) |
| *Streptococcus mitis (01)* | blood culture | clindamycin (0.19) |
| *Streptococcus oralis (01)* | blood culture | clindamycin (>256) |
| *Streptococcus oralis (02)* | blood culture | clindamycin (>256) |
| *Streptococcus pneumoniae (01)* | blood culture | erythromycin |
| *Streptococcus pneumoniae (02)* | blood culture | erythromycin |
| *Streptococcus pneumoniae (03)* | blood culture | erythromycin |
| *Streptococcus pneumoniae (04)* | blood culture | erythromycin |
| *Streptococcus pneumoniae (05)* | blood culture | erythromycin |
| *Streptococcus pneumoniae (06)* | blood culture | erythromycin |
| *Streptococcus pneumoniae (07)* | blood culture | erythromycin |
| *Streptococcus pneumoniae (08)* | blood culture | erythromycin |
| *Streptococcus pneumoniae (09)* | blood culture | erythromycin |
| *Streptococcus pneumoniae (10)* | blood culture | erythromycin |
| *Streptococcus pneumoniae (11)* | blood culture | erythromycin |
| *Streptococcus pneumoniae (12)* | blood culture | erythromycin |

*Supplementary Table 2: Overview of the 27 clinical bacterial isolates used in the study (UNIL = University of Lausanne, Lausanne Switzerland; SwissTPH = Swiss Tropical and Public Health Institute, Allschwil, Switzerland).*

| isolate | enrichment site | country of origin | sample type | enrichment media |
|---|---|---|---|---|
| *Blautia luti (01)* | UNIL | Lao PDR | stool | mGAM |
| *Clostridium baratii (01)* | UNIL | Lao PDR | stool | Schaedler |
| *Clostridium baratii (02)* | UNIL | Lao PDR | stool | mGAM |
| *Clostridium perfringens (01)* | UNIL | Lao PDR | stool | BHI + inulin |
| *Clostridium perfringens (02)* | UNIL | Lao PDR | stool | BHI + inulin |
| *Enterococcus faecalis (01)* | SwissTPH | Lao PDR | stool | TH |
| *Enterococcus faecalis (02)* | SwissTPH | Lao PDR | stool | LB |
| *Enterococcus faecalis (03)* | SwissTPH | Lao PDR | stool | TH |
| *Enterococcus faecium (01)* | SwissTPH | Lao PDR | stool | BHI + 5% yeast |
| *Enterococcus faecium (02)* | SwissTPH | Lao PDR | stool | LB |
| *Enterococcus faecium (03)* | SwissTPH | Lao PDR | stool | mGAM |
| *Enterococcus hirae (01)* | SwissTPH | Lao PDR | stool | BHI + 5% yeast |
| *Enterococcus hirae (02)* | SwissTPH | Lao PDR | stool | BHI + 5% yeast |
| *Enterococcus hirae (03)* | SwissTPH | Lao PDR | stool | BHI + 5% yeast |
| *Enterococcus hirae (04)* | SwissTPH | Lao PDR | stool | BHI + 5% yeast |
| *Enterococcus hirae (05)* | SwissTPH | Lao PDR | stool | BHI + 5% yeast |
| *Enterococcus hirae (06)* | SwissTPH | Lao PDR | stool | TH |
| *Escherichia coli (01)* | SwissTPH | Lao PDR | stool | LB |
| *Escherichia coli (02)* | SwissTPH | Lao PDR | stool | TH |

| | | | | |
|---|---|---|---|---|
| *Escherichia coli (03)* | UNIL | Lao PDR | stool | mGAM |
| *Escherichia coli (04)* | UNIL | Lao PDR | stool | BHI + inulin |
| *Escherichia coli (05)* | UNIL | Lao PDR | stool | Schaedler |
| *Streptococcus anginosus (01)* | SwissTPH | Lao PDR | stool | TH |
| *Streptococcus equinus (01)* | SwissTPH | Lao PDR | stool | BHI + 5% yeast |
| *Streptococcus equinus (02)* | SwissTPH | Lao PDR | stool | mGAM |
| *Streptococcus lutetiensis (01)* | SwissTPH | Lao PDR | stool | BHI + 5% yeast |
| *Streptococcus sanguinis(01)* | UNIL | Lao PDR | stool | mGAM |

*Supplementary Table 3: Overview of the 12 commercial bacterial isolates used in the study (UNIL = University of Lausanne, Lausanne Switzerland).*

| **isolate** | **strain designation** | **source** |
|---|---|---|
| *Bacteroides stercoris (01)* | DSM 19555 | Ordered |
| *Bacteroides uniformis (01)* | DSM 6597 | Ordered |
| *Blautia obeum (01)* | DSM 25238 | Provided by UNIL |
| *Blautia luti (02)* | DSM 19850 | Provided by UNIL |
| *Blautia luti (03)* | DSM 25403 | Provided by UNIL |
| *Dorea formicigenerans (01)* | JCM 31256 | Provided by UNIL |
| *Dorea formicigenerans (02)* | DSM 3992 | Ordered |
| *Dorea longicatena (01)* | DSM 13814 | Ordered |
| *Lactobacillus salivarius (01)* | DSM 20555 | Ordered |
| *Staphylococcus aureus (01)* | DSM 20231 | Ordered |
| *Streptococcus parasanguinis (01)* | DSM 6778 | Ordered |
| *Streptococcus salivarius (01)* | DSM 20067 | Ordered |

*Supplementary Table 4: Overview of the 10 challenged isolates used in the study. Final challenging concentrations are indicated in brackets.*

| **isolate** | **source** | **resistance** |
|---|---|---|
| *S. salivarius (01-IVc)* | *Streptococcus salivarius (01)*, see Supplementary Table 3 | ivermectin (20µM) |
| *S. salivarius (01-MXc)* | *Streptococcus salivarius (01)*, see Supplementary Table 3 | moxidectin (20µM) |
| *S. parasanguinis (01-IVc)* | *Streptococcus parasanguinis (01)*, see Supplementary Table 3 | ivermectin (20µM) |
| *S. parasanguinis (01-MXc)* | *Streptococcus parasanguinis (01)*, see Supplementary Table 3 | moxidectin (20µM) |
| *S. pneumoniae (02-IVc)* | *Streptococcus pneumoniae (02)*, see Supplementary Table 1 | ivermectin (20µM) |
| *S. pneumoniae (02-MXc)* | *Streptococcus pneumoniae (02)*, see Supplementary Table 1 | moxidectin (10µM) |
| *S. mitis (01-IVc)* | *Streptococcus mitis (01)*, see Supplementary Table 1 | ivermectin (20µM) |
| *S. mitis (01-MXc)* | *Streptococcus mitis (01)*, see Supplementary Table 1 | moxidectin (10µM) |
| *S. dysgalactiae (01-IVc)* | *Streptococcus dysgalactiae (01)*, see Supplementary Table 1 | ivermectin (20µM) |
| *S. dysgalactiae (01-MXc)* | *Streptococcus dysgalactiae (01)*, see Supplementary Table 1 | moxidectin (20µM) |

*Supplementary Table 5: Pairwise Wilcoxon rank sum test of AUC ratios for 11 co-incubations in presence of anthelminthics (IV = ivermectin, MX = moxidectin) and antibiotics (EM = erythromycin, CH = clarithromycin, AZ = azithromycin, CL = clindamycin, CX = ciprofloxacin, TC = tetracycline, MP = meropenem, IP = imipenem) at 1µM.*

**Pairwise Wilcoxon rank sum test (1µM)**

| | AZ | CH | CL | CX | EM | IP | IV | MP | MX |
|---|---|---|---|---|---|---|---|---|---|
| **CH** | 0.086 | - | | | | | | | |
| **CL** | 0.147 | 0.918 | - | | | | | | |
| **CX** | 0.000 | 0.000 | 0.000 | - | | | | | |
| **EM** | 0.329 | 0.248 | 0.393 | 0.000 | - | | | | |
| **IP** | 0.031 | 0.076 | 0.159 | 0.000 | 0.035 | - | | | |
| **IV** | 0.000 | 0.000 | 0.000 | 0.749 | 0.000 | 0.001 | - | | |
| **MP** | 0.035 | 0.114 | 0.303 | 0.001 | 0.067 | 0.886 | 0.001 | - | |
| **MX** | 0.000 | 0.000 | 0.000 | 0.632 | 0.000 | 0.000 | 0.665 | 0.001 | - |
| **TC** | 0.949 | 0.248 | 0.159 | 0.006 | 0.463 | 0.031 | 0.006 | 0.049 | 0.011 |

*Supplementary Table 6: Pairwise Wilcoxon rank sum test of AUC ratios for 11 co-incubations in presence of anthelminthics (IV = ivermectin, MX = moxidectin) and antibiotics (EM = erythromycin, CH = clarithromycin, AZ = azithromycin, CL = clindamycin, CX = ciprofloxacin, TC = tetracycline, MP = meropenem, IP = imipenem) at 5µM.*

**Pairwise Wilcoxon rank sum test (5µM)**

| | AZ | CH | CL | CX | EM | IP | IV | MP | MX |
|---|---|---|---|---|---|---|---|---|---|
| **CH** | 0.025 | - | | | | | | | |
| **CL** | 0.531 | 0.409 | - | | | | | | |
| **CX** | 0.030 | 0.005 | 0.014 | - | | | | | |
| **EM** | 0.249 | 0.290 | 0.787 | 0.005 | - | | | | |
| **IP** | 0.005 | 0.187 | 0.086 | 0.004 | 0.025 | - | | | |
| **IV** | 0.753 | 0.066 | 0.235 | 0.187 | 0.183 | 0.014 | - | | |
| **MP** | 0.011 | 0.380 | 0.149 | 0.004 | 0.086 | 0.970 | 0.025 | - | |
| **MX** | 0.939 | 0.086 | 0.667 | 0.086 | 0.165 | 0.025 | 0.875 | 0.029 | - |
| **TC** | 0.908 | 0.352 | 0.718 | 0.187 | 0.702 | 0.066 | 0.718 | 0.086 | 1.000 |

*Supplementary Table 7: Pairwise Wilcoxon rank sum test of AUC ratios for 11 co-incubations in presence of anthelminthics (IV = ivermectin, MX = moxidectin) and antibiotics (EM = erythromycin, CH = clarithromycin, AZ = azithromycin, CL = clindamycin, CX = ciprofloxacin, TC = tetracycline, MP = meropenem, IP = imipenem) at 10µM.*

**Pairwise Wilcoxon rank sum test (10µM)**

| | AZ | CH | CL | CX | EM | IP | IV | MP | MX |
|---|---|---|---|---|---|---|---|---|---|
| **CH** | 0.161 | - | | | | | | | |
| **CL** | 0.870 | 0.266 | - | | | | | | |
| **CX** | 0.632 | 0.113 | 0.477 | - | | | | | |
| **EM** | 0.287 | 0.584 | 0.619 | 0.220 | - | | | | |
| **IP** | 0.035 | 0.657 | 0.220 | 0.035 | 0.266 | - | | | |
| **IV** | 0.657 | 0.550 | 0.904 | 0.517 | 0.657 | 0.310 | - | | |
| **MP** | 0.035 | 0.657 | 0.220 | 0.035 | 0.266 | 0.949 | 0.266 | - | |
| **MX** | 0.266 | 0.949 | 0.517 | 0.220 | 0.657 | 0.657 | 0.657 | 0.657 | - |
| **TC** | 0.657 | 0.508 | 0.745 | 0.632 | 0.870 | 0.266 | 0.937 | 0.266 | 0.657 |

*Supplementary Table 8: Pearson correlation matrix of AUC ratios for 11 co-incubations in presence of anthelminthics (IV = ivermectin, MX = moxidectin) and macrolide/lincosamide antibiotics (EM = erythromycin, CH = clarithromycin, AZ = azithromycin, CL = clindamycin) at 1µM. Values in bold are different from 0 with a significance level alpha=0.05.*

| Variables | IV | MX | EM | CH | AZ | CL |
|---|---|---|---|---|---|---|
| IV | **1** | **0.747** | 0.349 | 0.493 | 0.410 | 0.096 |
| MX | **0.747** | **1** | 0.518 | **0.756** | 0.575 | 0.088 |
| EM | 0.349 | 0.518 | **1** | **0.920** | **0.954** | 0.359 |
| CH | 0.493 | **0.756** | **0.920** | **1** | **0.926** | 0.203 |
| AZ | 0.410 | 0.575 | **0.954** | **0.926** | **1** | 0.207 |
| CL | 0.096 | 0.088 | 0.359 | 0.203 | 0.207 | **1** |

*Supplementary Table 9: Pearson correlation p-values of AUC ratios for 11 co-incubations in presence of anthelminthics (IV = ivermectin, MX = moxidectin) and macrolide/lincosamide antibiotics (EM = erythromycin, CH = clarithromycin, AZ = azithromycin, CL = clindamycin) at 1µM.*

| Variables | IV | MX | EM | CH | AZ | CL |
|---|---|---|---|---|---|---|
| IV | **0** | **0.008** | 0.293 | 0.124 | 0.210 | 0.779 |
| MX | **0.008** | **0** | 0.103 | **0.007** | 0.064 | 0.798 |
| EM | 0.293 | 0.103 | **0** | **<0.0001** | **<0.0001** | 0.278 |
| CH | 0.124 | **0.007** | **<0.0001** | **0** | **<0.0001** | 0.549 |
| AZ | 0.210 | 0.064 | **<0.0001** | **<0.0001** | **0** | 0.541 |
| CL | 0.779 | 0.798 | 0.278 | 0.549 | 0.541 | **0** |

*Supplementary Table 10: Pearson correlation matrix of AUC ratios for 11 co-incubations in presence of anthelminthics (IV = ivermectin, MX = moxidectin) and macrolide/lincosamide antibiotics (EM = erythromycin, CH = clarithromycin, AZ = azithromycin, CL = clindamycin) at 5µM. Values in bold are different from 0 with a significance level alpha=0.05.*

| Variables | IV | MX | EM | CH | AZ | CL |
|---|---|---|---|---|---|---|
| IV | **1** | 0.322 | **0.647** | 0.590 | **0.826** | 0.150 |
| MX | 0.322 | **1** | **0.608** | 0.595 | 0.285 | 0.337 |
| EM | **0.647** | **0.608** | **1** | **0.957** | **0.792** | 0.426 |
| CH | 0.590 | 0.595 | **0.957** | **1** | **0.639** | 0.381 |
| AZ | **0.826** | 0.285 | **0.792** | **0.639** | **1** | 0.163 |
| CL | 0.150 | 0.337 | 0.426 | 0.381 | 0.163 | **1** |

*Supplementary Table 11: Pearson correlation p-values of AUC ratios for 11 co-incubations in presence of anthelminthics (IV = ivermectin, MX = moxidectin) and macrolide/lincosamide antibiotics (EM = erythromycin, CH = clarithromycin, AZ = azithromycin, CL = clindamycin) at 5µM.*

| Variables | IV | MX | EM | CH | AZ | CL |
|---|---|---|---|---|---|---|
| IV | **0** | 0.334 | **0.032** | 0.056 | **0.002** | 0.660 |
| MX | 0.334 | **0** | **0.047** | 0.053 | 0.396 | 0.310 |
| EM | **0.032** | **0.047** | **0** | **<0.0001** | **0.004** | 0.192 |
| CH | 0.056 | 0.053 | **<0.0001** | **0** | **0.034** | 0.247 |
| AZ | **0.002** | 0.396 | **0.004** | **0.034** | **0** | 0.632 |
| CL | 0.660 | 0.310 | 0.192 | 0.247 | 0.632 | **0** |

*Supplementary Table 12: Pearson correlation matrix of AUC ratios for 11 co-incubations in presence of anthelminthics (IV = ivermectin, MX = moxidectin) and macrolide/lincosamide antibiotics (EM = erythromycin, CH = clarithromycin, AZ = azithromycin, CL = clindamycin) at 10µM. Values in bold are different from 0 with a significance level alpha=0.05.*

| Variables | IV | MX | EM | CH | AZ | CL |
|---|---|---|---|---|---|---|
| IV | **1** | 0.170 | **0.651** | **0.731** | **0.706** | 0.155 |
| MX | 0.170 | **1** | 0.475 | 0.529 | 0.231 | 0.253 |
| EM | **0.651** | 0.475 | **1** | **0.923** | **0.913** | 0.449 |
| CH | **0.731** | 0.529 | **0.923** | **1** | **0.800** | 0.363 |
| AZ | **0.706** | 0.231 | **0.913** | **0.800** | **1** | 0.426 |
| CL | 0.155 | 0.253 | 0.449 | 0.363 | 0.426 | **1** |

*Supplementary Table 13: Pearson correlation p-values of AUC ratios for 11 co-incubations in presence of anthelminthics (IV = ivermectin, MX = moxidectin) and macrolide/lincosamide antibiotics (EM = erythromycin, CH = clarithromycin, AZ = azithromycin, CL = clindamycin) at 10µM.*

| Variables | IV | MX | EM | CH | AZ | CL |
|---|---|---|---|---|---|---|
| IV | **0** | 0.618 | **0.030** | **0.011** | **0.015** | 0.650 |
| MX | 0.618 | **0** | 0.140 | 0.094 | 0.494 | 0.452 |
| EM | **0.030** | 0.140 | **0** | **<0.0001** | **<0.0001** | 0.166 |
| CH | **0.011** | 0.094 | **<0.0001** | **0** | **0.003** | 0.272 |
| AZ | **0.015** | 0.494 | **<0.0001** | **0.003** | **0** | 0.191 |
| CL | 0.650 | 0.452 | 0.166 | 0.272 | 0.191 | **0** |

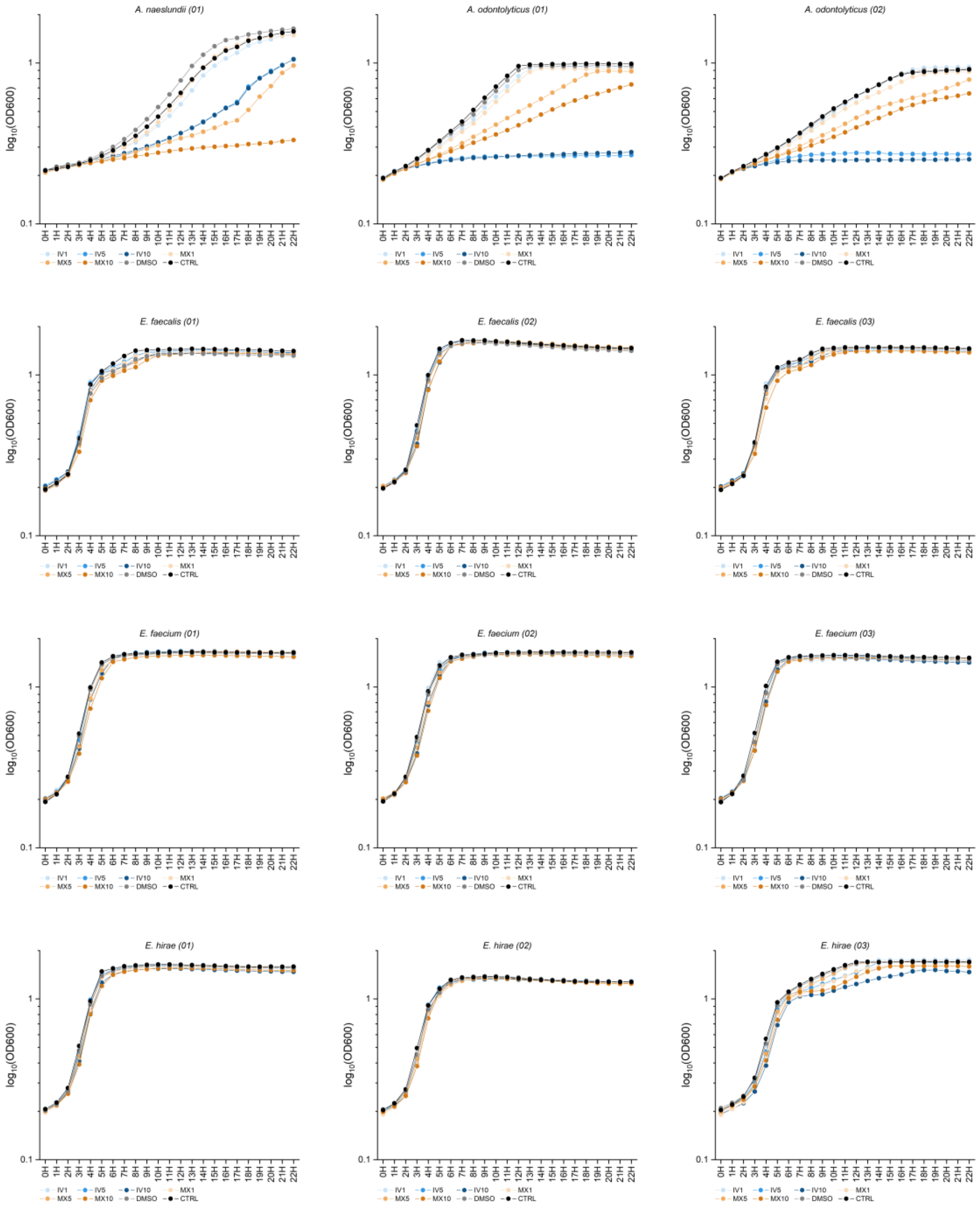


*Supplementary Figure 1: Aerobic growth curves of 12 bacterial isolates co-incubated with IV (blue) or MX (orange) at concentrations of either 1µM, 5µM or 10µM. Each datapoint represents an average of two independent experiments. The grey curves represent growth in presence of 0.2% DMSO, the black curve represents a positive growth control in presence of BHI + 5% yeast only. Left to right and top to bottom: A. naeslundii (01), A. odontolyticus (01), A. odontolyticus (02), E. faecalis (01), E. faecalis (02), E. faecalis (03), E. faecium (01), E. faecium (02), E. faecium (03), E. hirae (01), E. hirae (02) and E. hirae (03).*

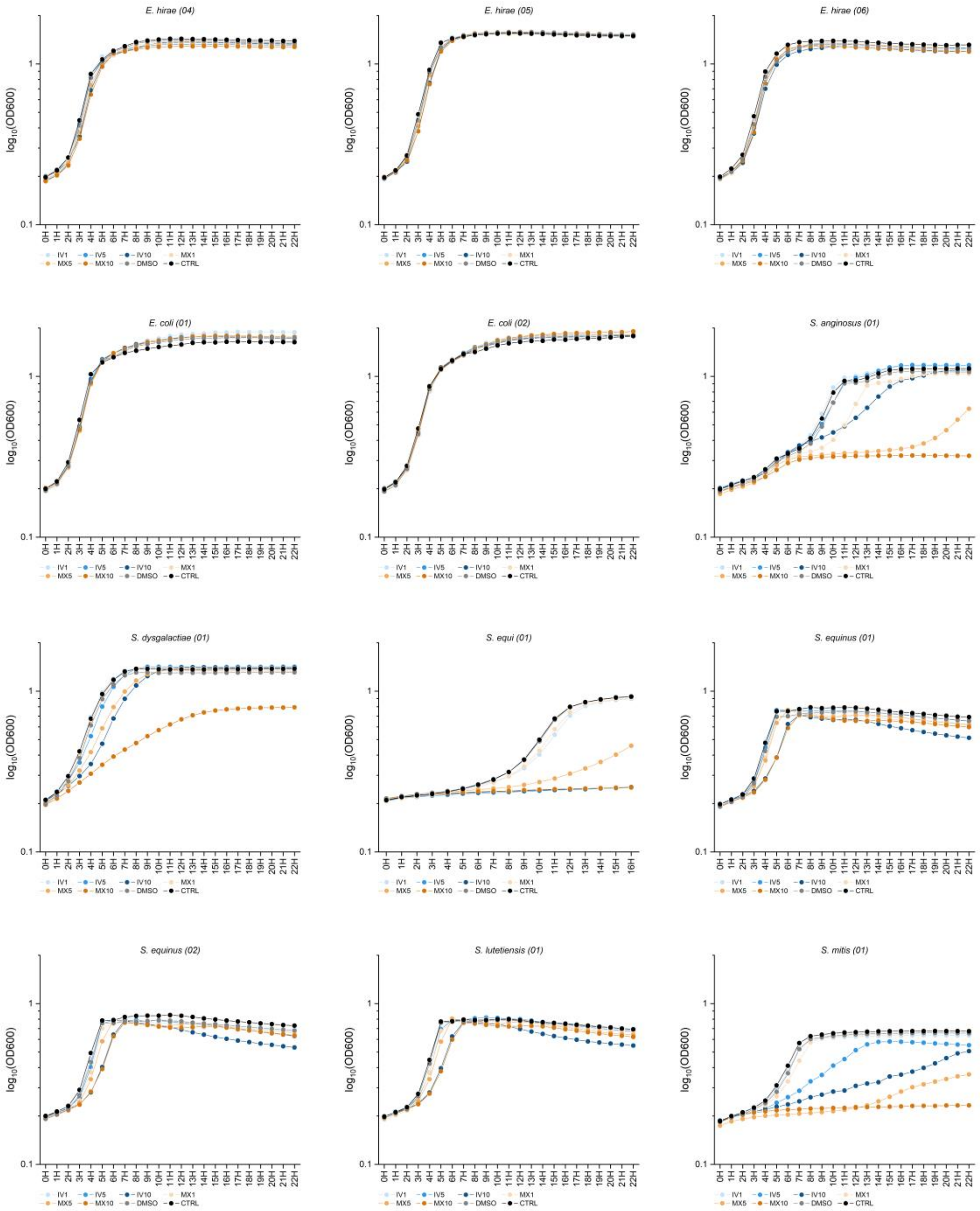


*Supplementary Figure 2: Aerobic growth curves of 12 bacterial isolates co-incubated with IV (blue) or MX (orange) at concentrations of either 1µM, 5µM or 10µM. Each datapoint represents an average of two independent experiments. The grey curves represent growth in presence of 0.2% DMSO, the black curve represents a positive growth control in presence of BHI + 5% yeast only. Left to right and top to bottom: E. hirae (04), E. hirae (05), E. hirae (06), E. coli (01), E. coli (02), S. anginosus (01), S. dysgalactiae (01), S. equi (01), S. equinus (01), S. equinus (02), S. lutetiensis (01) and S. mitis (01).*

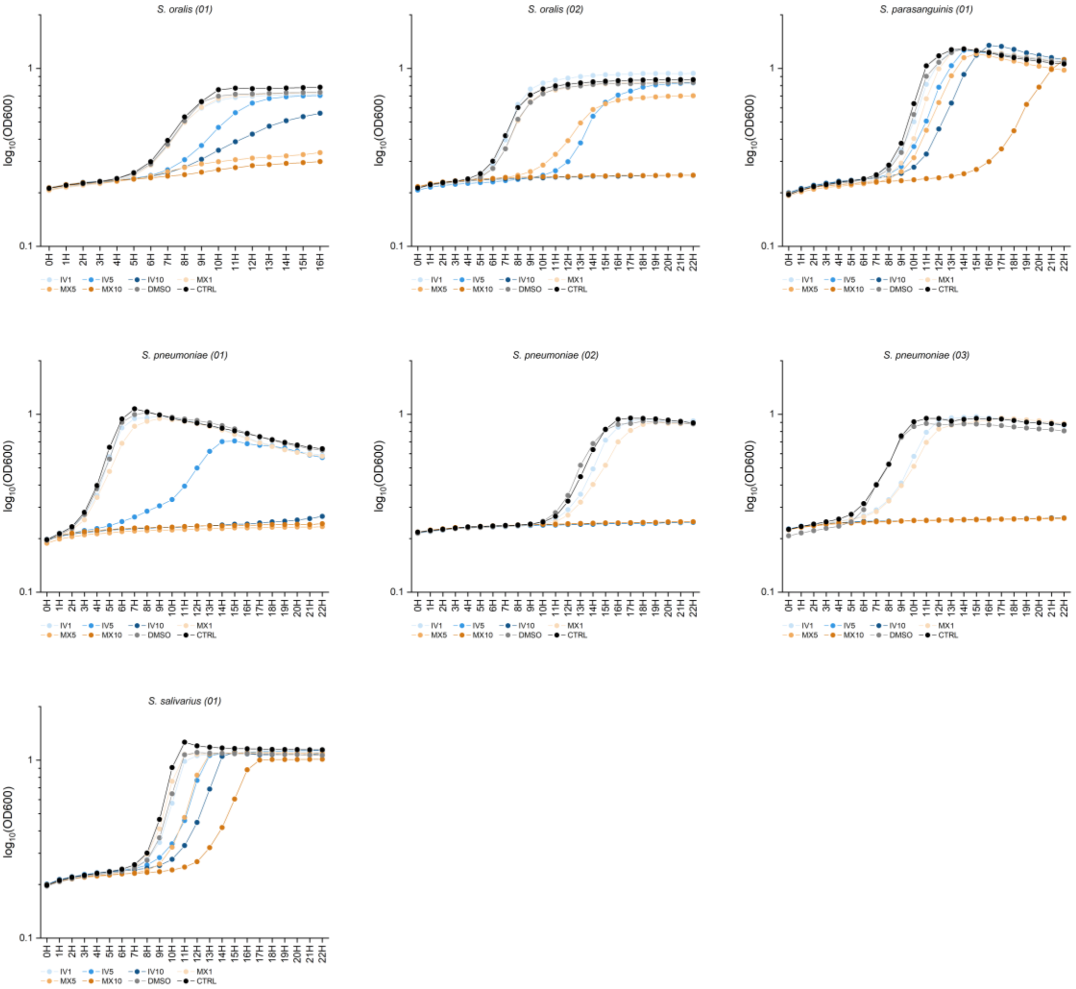


*Supplementary Figure 3: Aerobic growth curves of 7 bacterial isolates co-incubated with IV (blue) or MX (orange) at concentrations of either 1µM, 5µM or 10µM. Each datapoint represents an average of two independent experiments. The grey curves represent growth in presence of 0.2% DMSO, the black curve represents a positive growth control in presence of BHI + 5% yeast only. Left to right and top to bottom: S. oralis (01), S. oralis (02), S. parasanguinis (01), S. pneumoniae (01), S. pneumoniae (02), S. pneumoniae (03) and S. salivarius (01).*

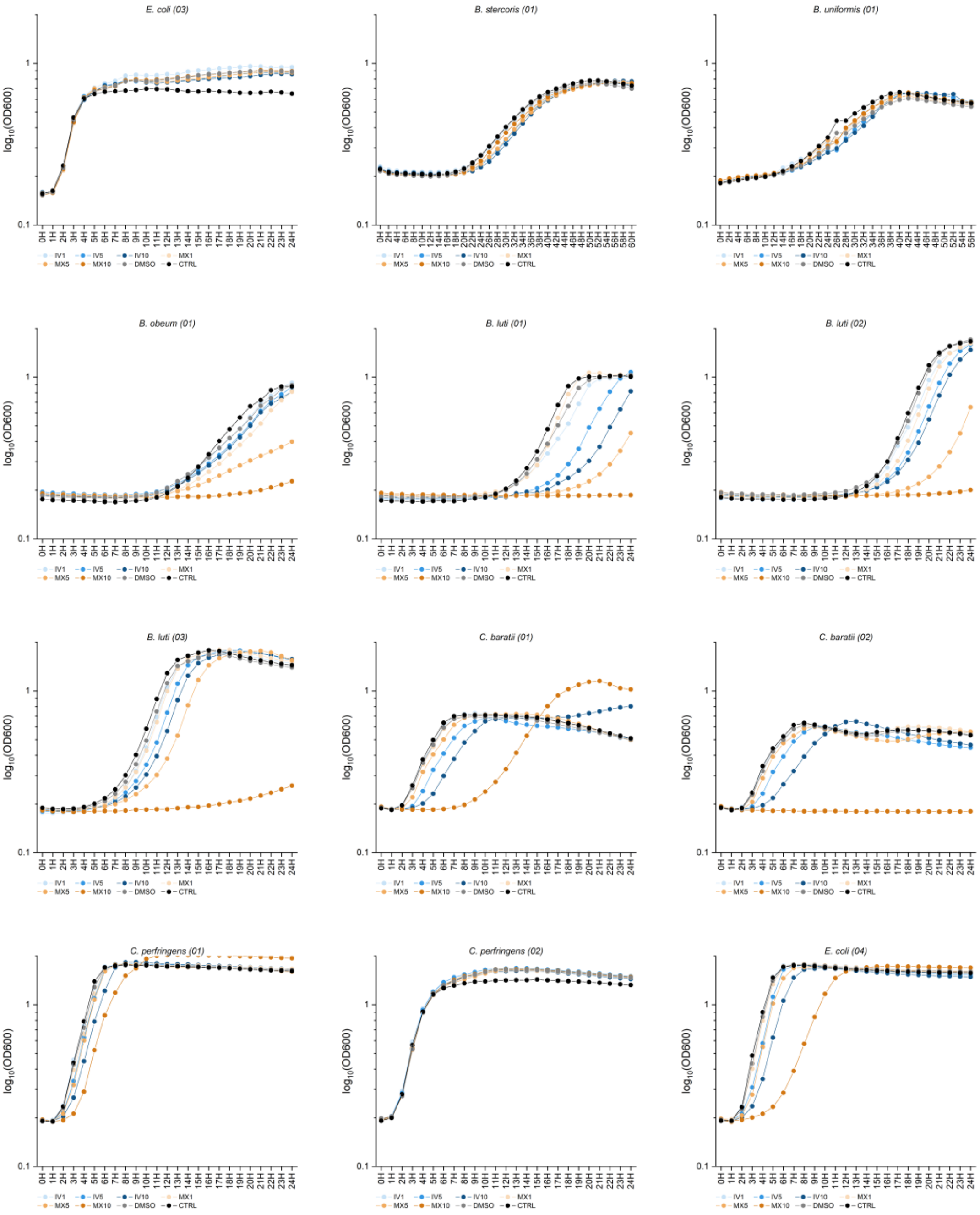


*Supplementary Figure 4: Anaerobic growth curves of 12 bacterial isolates co-incubated with IV (blue) or MX (orange) at concentrations of either 1µM, 5µM or 10µM. Each datapoint represents an average of two independent experiments. The grey curves represent growth in presence of 0.2% DMSO, the black curve represents a positive growth control in presence of BHI + 5% yeast only. Left to right and top to bottom: E. coli (03), B. stercoris (01), B. uniformis (01), B. obeum (01), B. luti (01), B. luti (02), B. luti (03), C. baratii (01), C. baratii (02), C. perfringens (01), C. perfringens (02) and E. coli (04).*

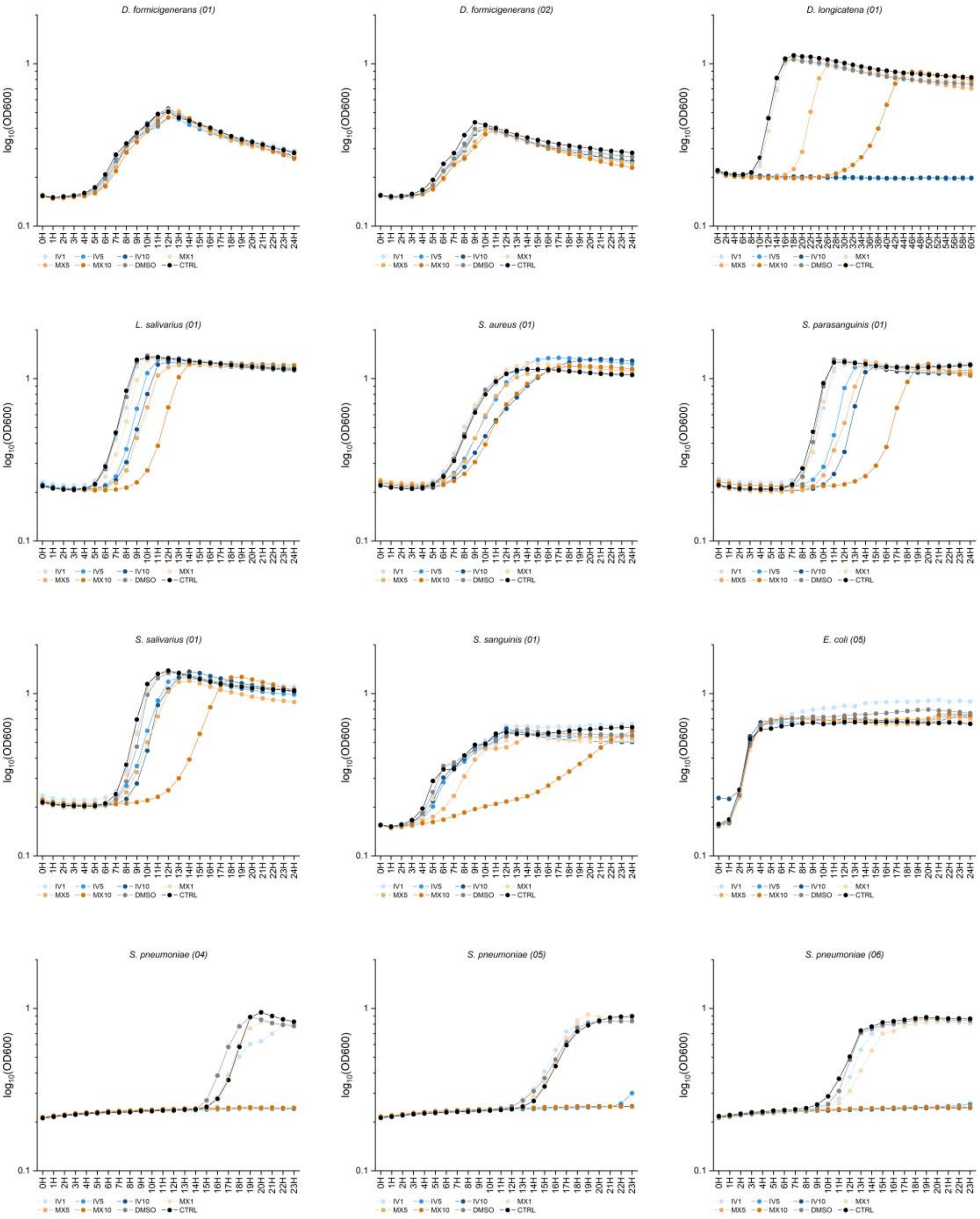


*Supplementary Figure 5: Anaerobic growth curves of 12 bacterial isolates co-incubated with IV (blue) or MX (orange) at concentrations of either 1µM, 5µM or 10µM. Each datapoint represents an average of two independent experiments. The grey curves represent growth in presence of 0.2% DMSO, the black curve represents a positive growth control in presence of BHI + 5% yeast only. Left to right and top to bottom: D. formicigenerans (01), D. formicigenerans (02), D. longicatena (01), L. salivarius (01), S. aureus (01), S. parasanguinis (01), S. salivarius (01), S. sanguinis (01), E. coli (05), S. pneumoniae (04), S. pneumoniae (05) and S. pneumoniae (06).*

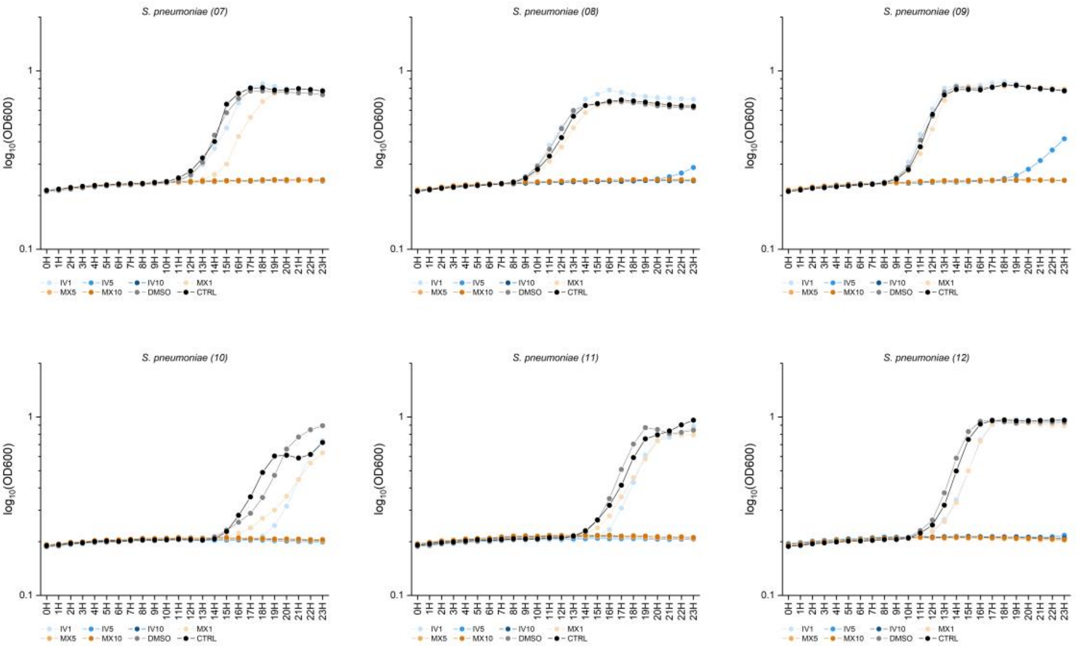


*Supplementary Figure 6: Anaerobic growth curves of 6 bacterial isolates co-incubated with IV (blue) or MX (orange) at concentrations of either 1µM, 5µM or 10µM. Each datapoint represents an average of two independent experiments. The grey curves represent growth in presence of 0.2% DMSO, the black curve represents a positive growth control in presence of BHI + 5% yeast only. Left to right and top to bottom: S. pneumoniae (07), S. pneumoniae (08), S. pneumoniae (09), S. pneumoniae (10), S. pneumoniae (11) and S. pneumoniae (12).*

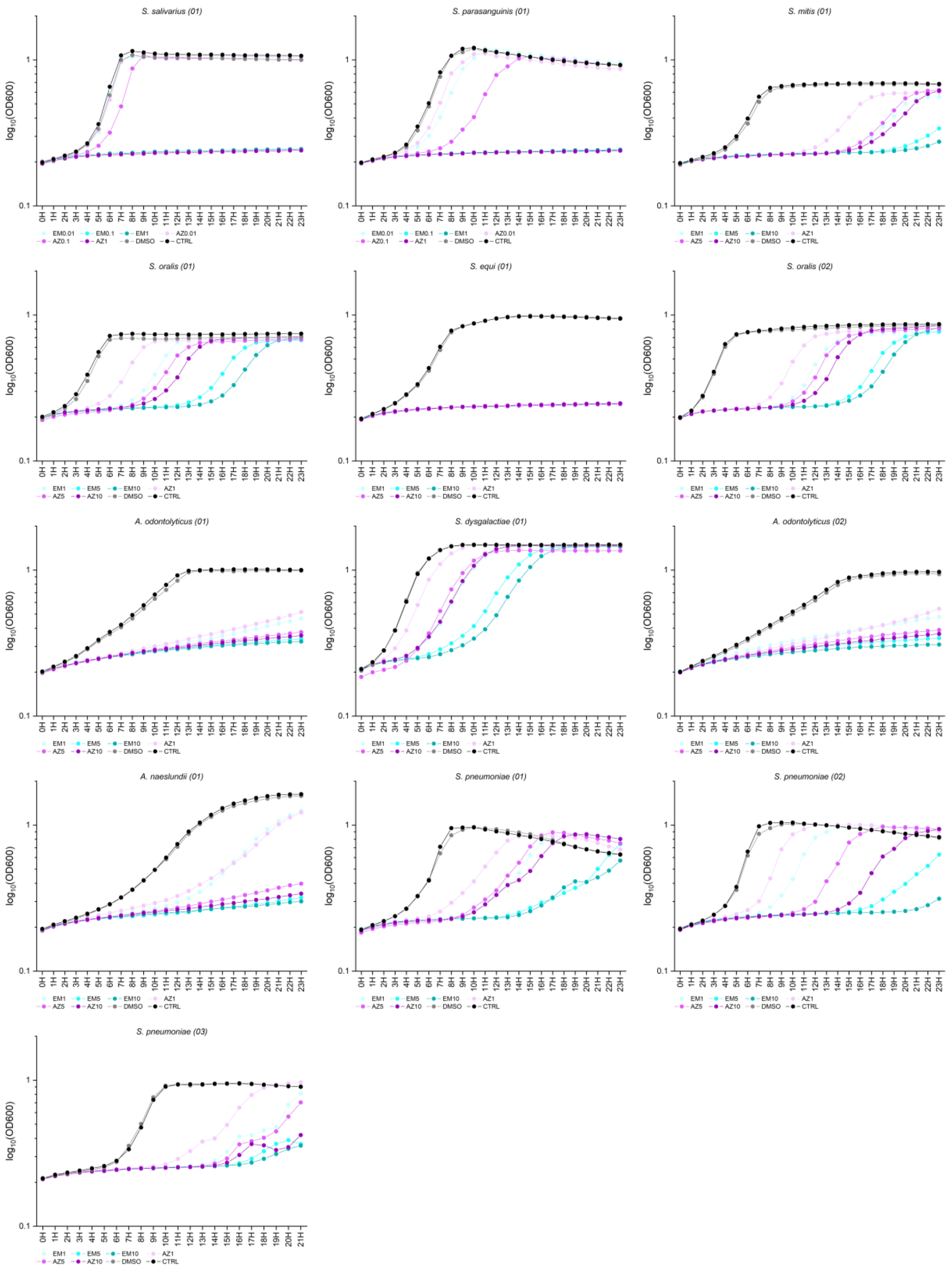


*Supplementary Figure 7: Aerobic growth curves of 13 bacterial isolates co-incubated with EM (turquoise) or AZ (purple) at three different concentrations. Each datapoint represents an average of two independent experiments. The grey curves represent growth in presence of 0.2% DMSO, the black curve represents a positive growth control in presence of BHI + 5% yeast only. Left to right and top to bottom: S. salivarius (01), S. parasanguinis (01), S. mitis (01), S. oralis (01), S. equi (01), S. oralis (02), A. odontolyticus (01), S. dysgalactiae (01), A. odontolyticus (02), A. naeslundii (01), S. pneumoniae (01), S. pneumoniae (02) and S. pneumoniae (03).*

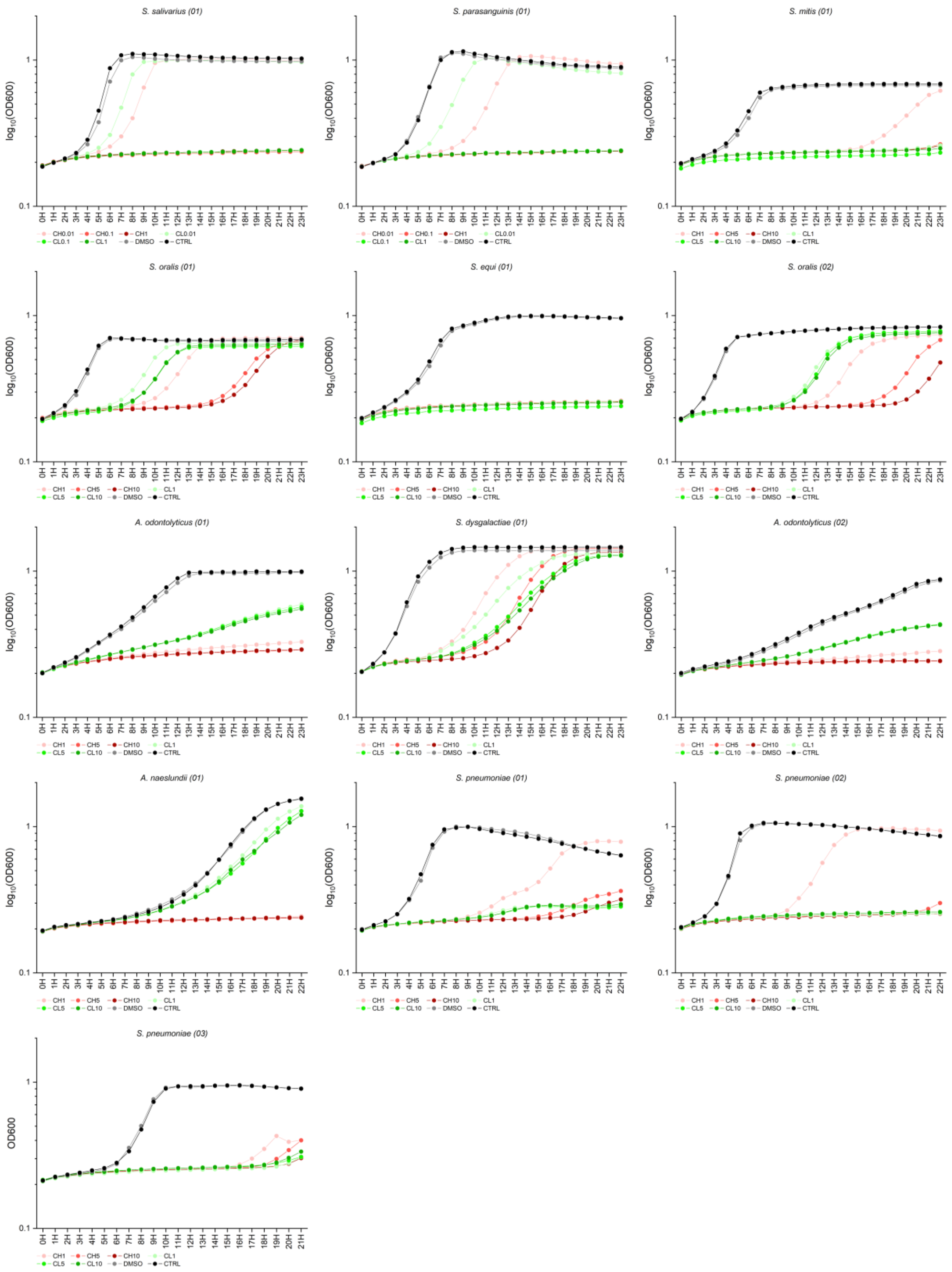


*Supplementary Figure 8: Aerobic growth curves of 13 bacterial isolates co-incubated with CH (red) or CL (green) at three different concentrations. Each datapoint represents an average of two independent experiments. The grey curves represent growth in presence of 0.2% DMSO, the black curve represents a positive growth control in presence of BHI + 5% yeast only. Left to right and top to bottom: S. salivarius (01), S. parasanguinis (01), S. mitis (01), S. oralis (01), S. equi (01), S. oralis (02), A. odontolyticus (01), S. dysgalactiae (01), A. odontolyticus (02), A. naeslundii (01), S. pneumoniae (01), S. pneumoniae (02) and S. pneumoniae (03).*

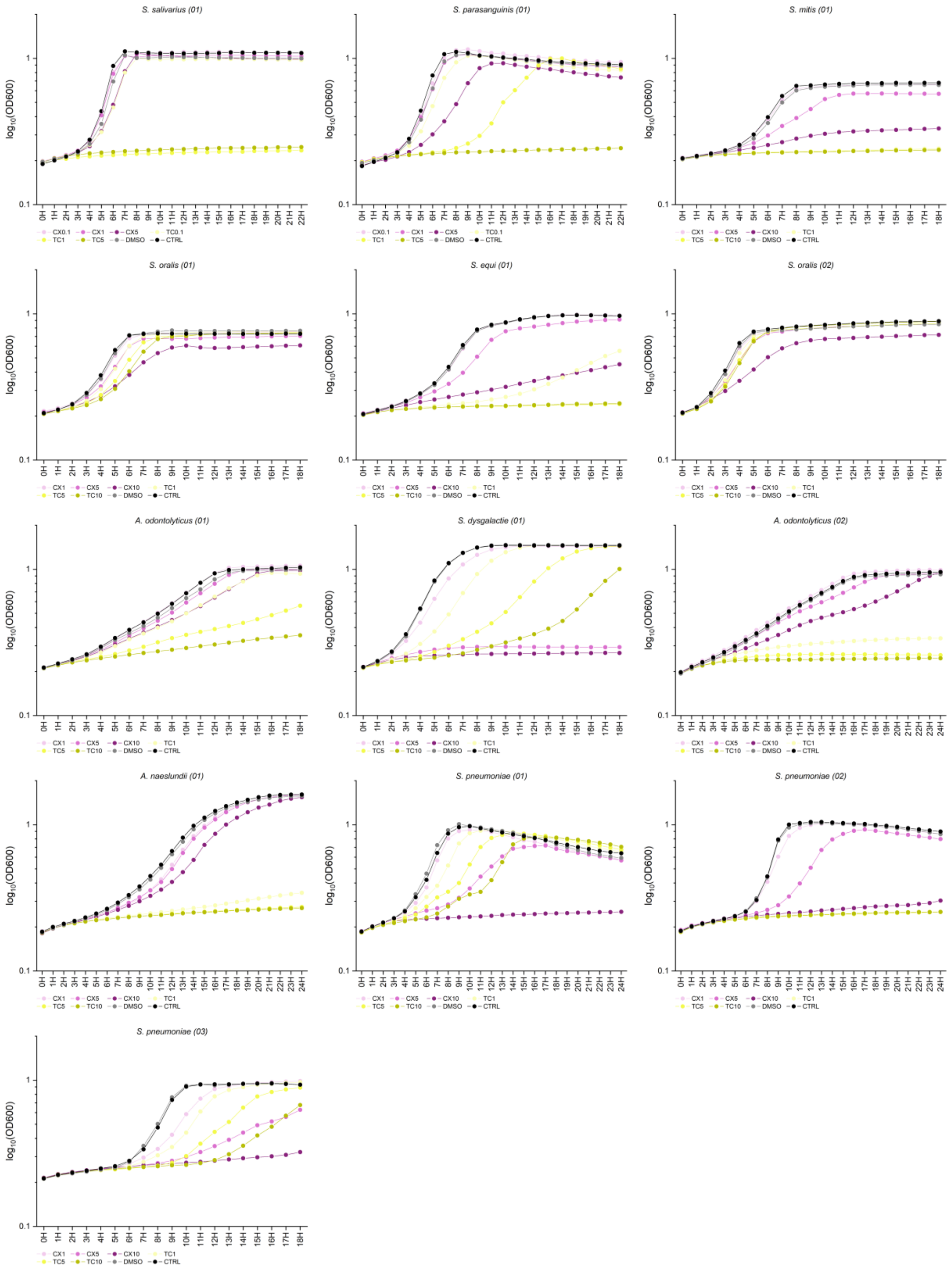


*Supplementary Figure 9: Aerobic growth curves of 13 bacterial isolates co-incubated with CX (mauve) or TC (yellow) at three different concentrations. Each datapoint represents an average of two independent experiments. The grey curves represent growth in presence of 0.2% DMSO, the black curve represents a positive growth control in presence of BHI + 5% yeast only. Left to right and top to bottom: S. salivarius (01), S. parasanguinis (01), S. mitis (01), S. oralis (01), S. equi (01), S. oralis (02), A. odontolyticus (01), S. dysgalactiae (01), A. odontolyticus (02), A. naeslundii (01), S. pneumoniae (01), S. pneumoniae (02) and S. pneumoniae (03).*

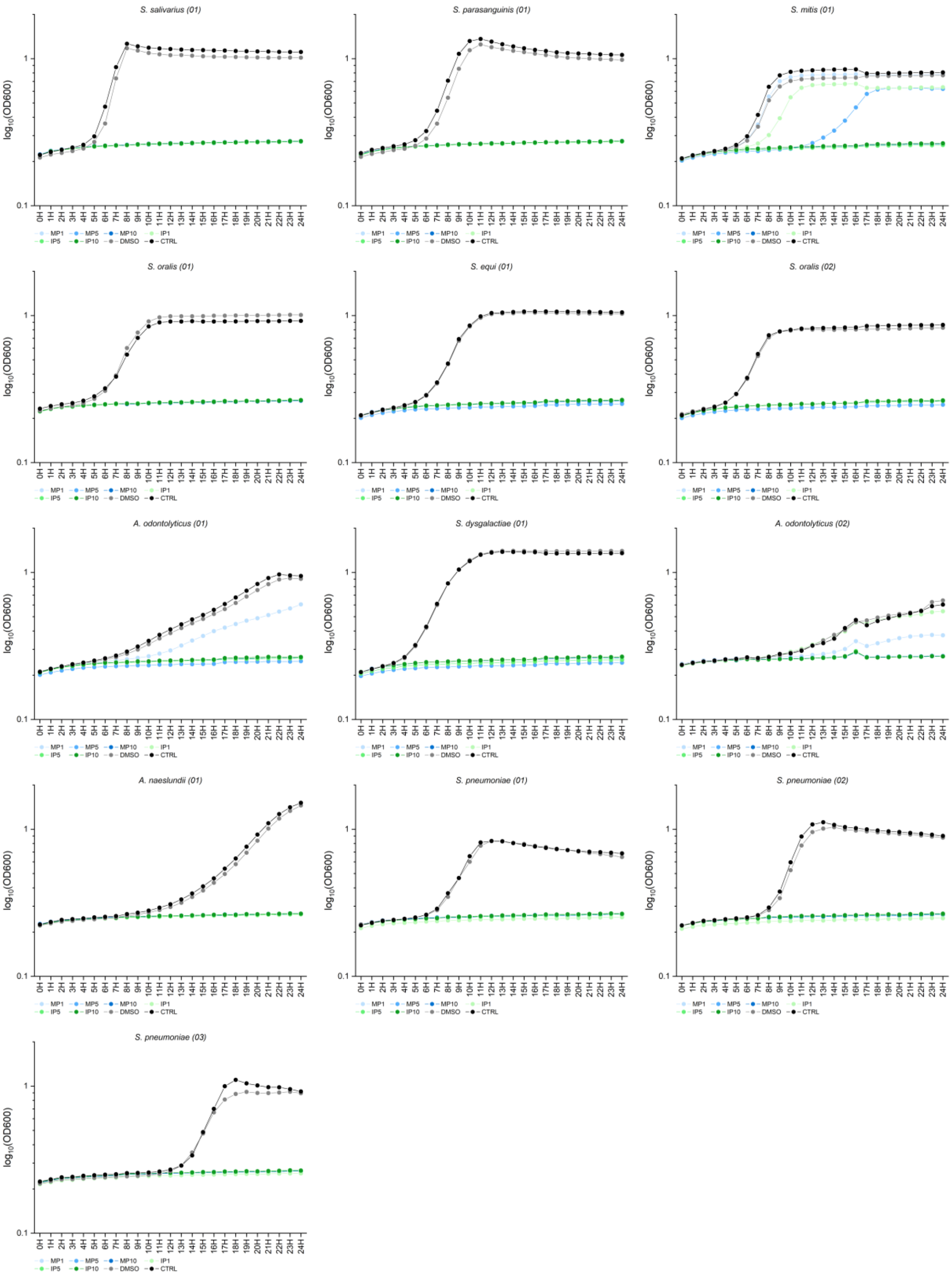


*Supplementary Figure 10: Aerobic growth curves of 13 bacterial isolates co-incubated with MP (blue) or IP (green) at three different concentrations. Each datapoint represents an average of two independent experiments. The grey curves represent growth in presence of 0.2% DMSO, the black curve represents a positive growth control in presence of BHI + 5% yeast only. Left to right and top to bottom: S. salivarius (01), S. parasanguinis (01), S. mitis (01), S. oralis (01), S. equi (01), S. oralis (02), A. odontolyticus (01), S. dysgalactiae (01), A. odontolyticus (02), A. naeslundii (01), S. pneumoniae (01), S. pneumoniae (02) and S. pneumoniae (03).*

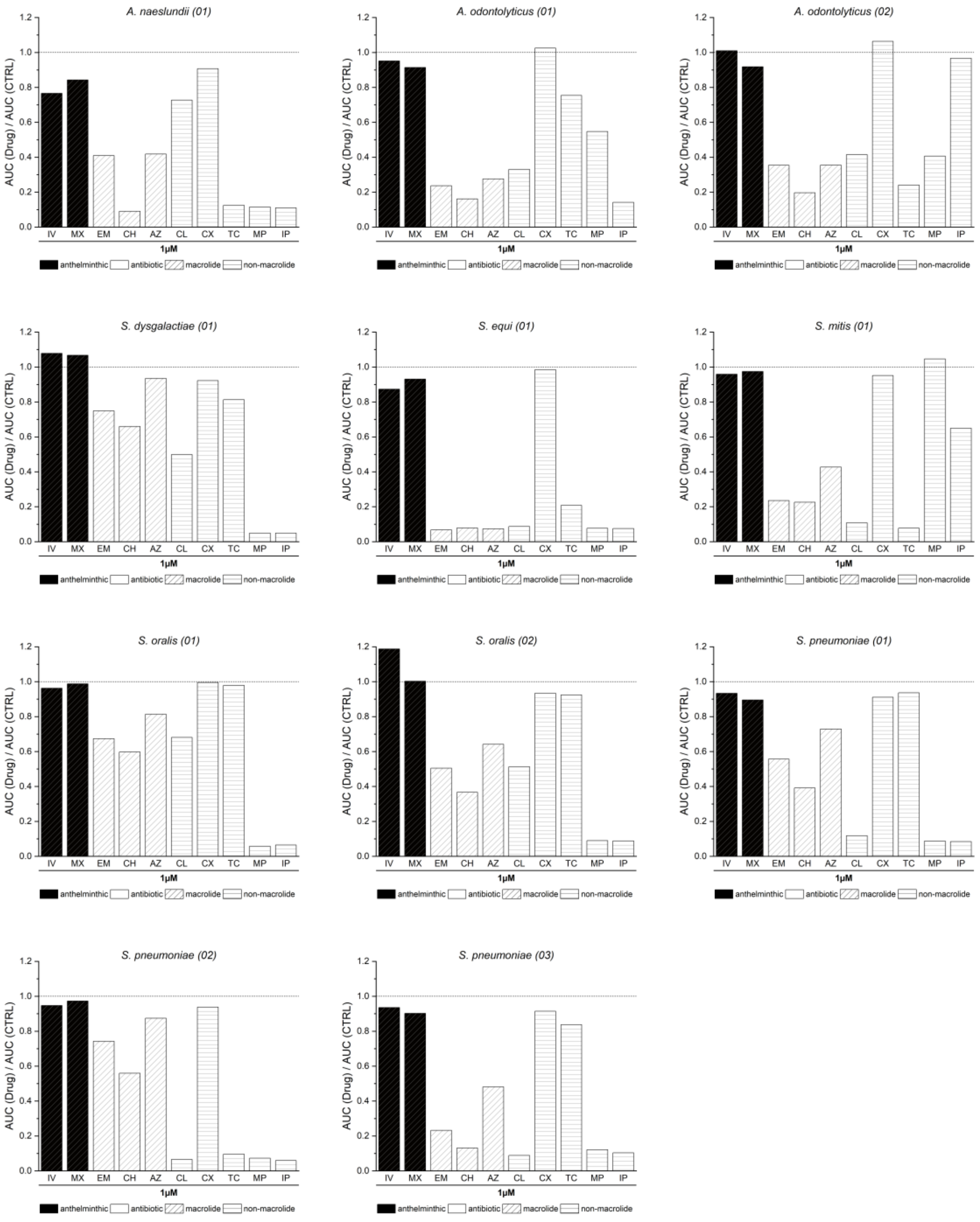


*Supplementary Figure 11: AUC ratios of 11 bacterial isolates in presence of IV, MX, EM, CH, AZ, CL, CX, TC, MP or IP at 1µM. The dotted line marks theoretical uninhibited growth of the isolate (AUC ratio = 1). Bar colors correspond to the primary usecase of the compound. Black = anthelminthic, white = antibiotic.*

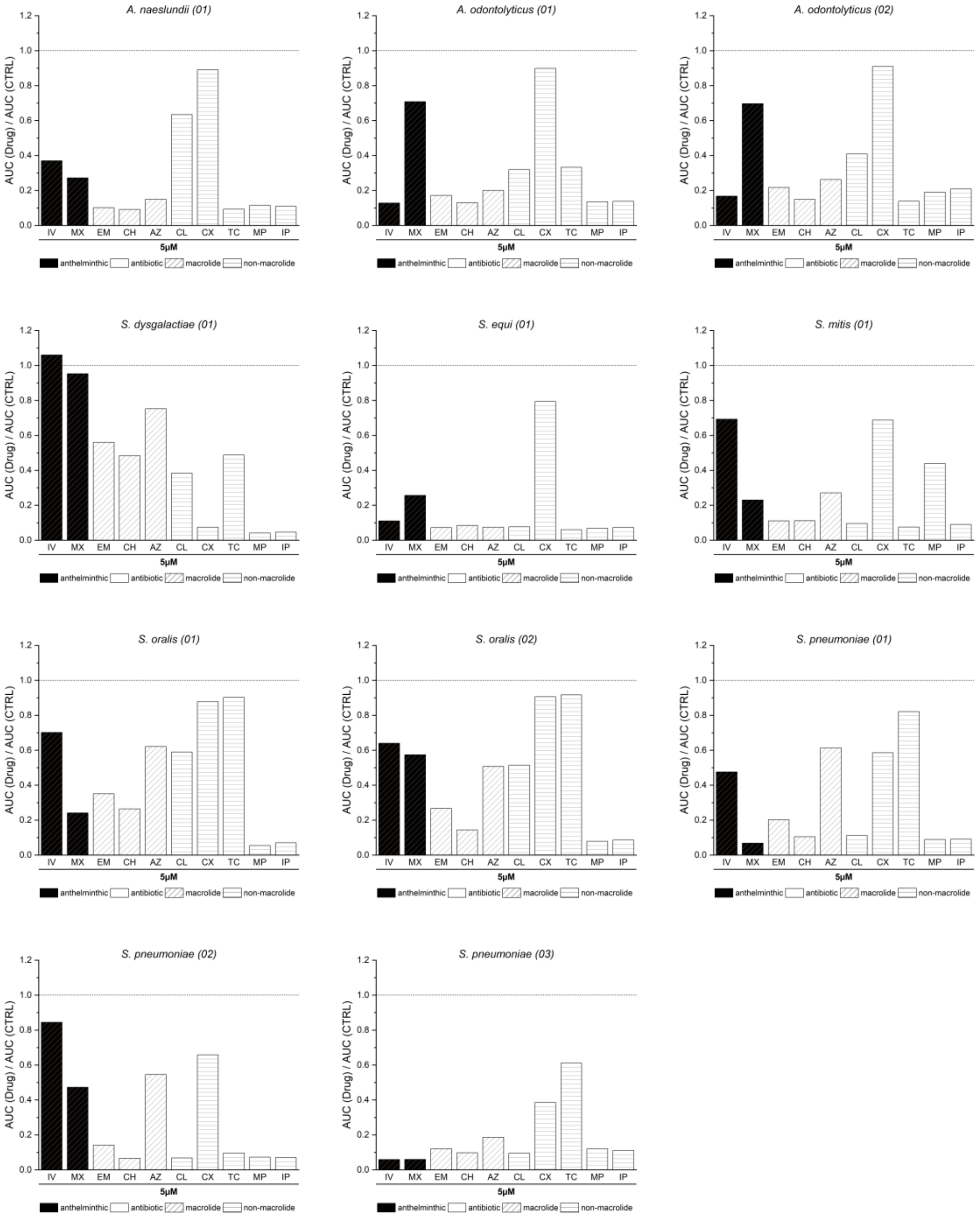


*Supplementary Figure 12: AUC ratios of 11 bacterial isolates in presence of IV, MX, EM, CH, AZ, CL, CX, TC, MP or IP at 5µM. The dotted line marks theoretical uninhibited growth of the isolate (AUC ratio = 1). Bar colors correspond to the primary usecase of the compound. Black = anthelminthic, white = antibiotic.*

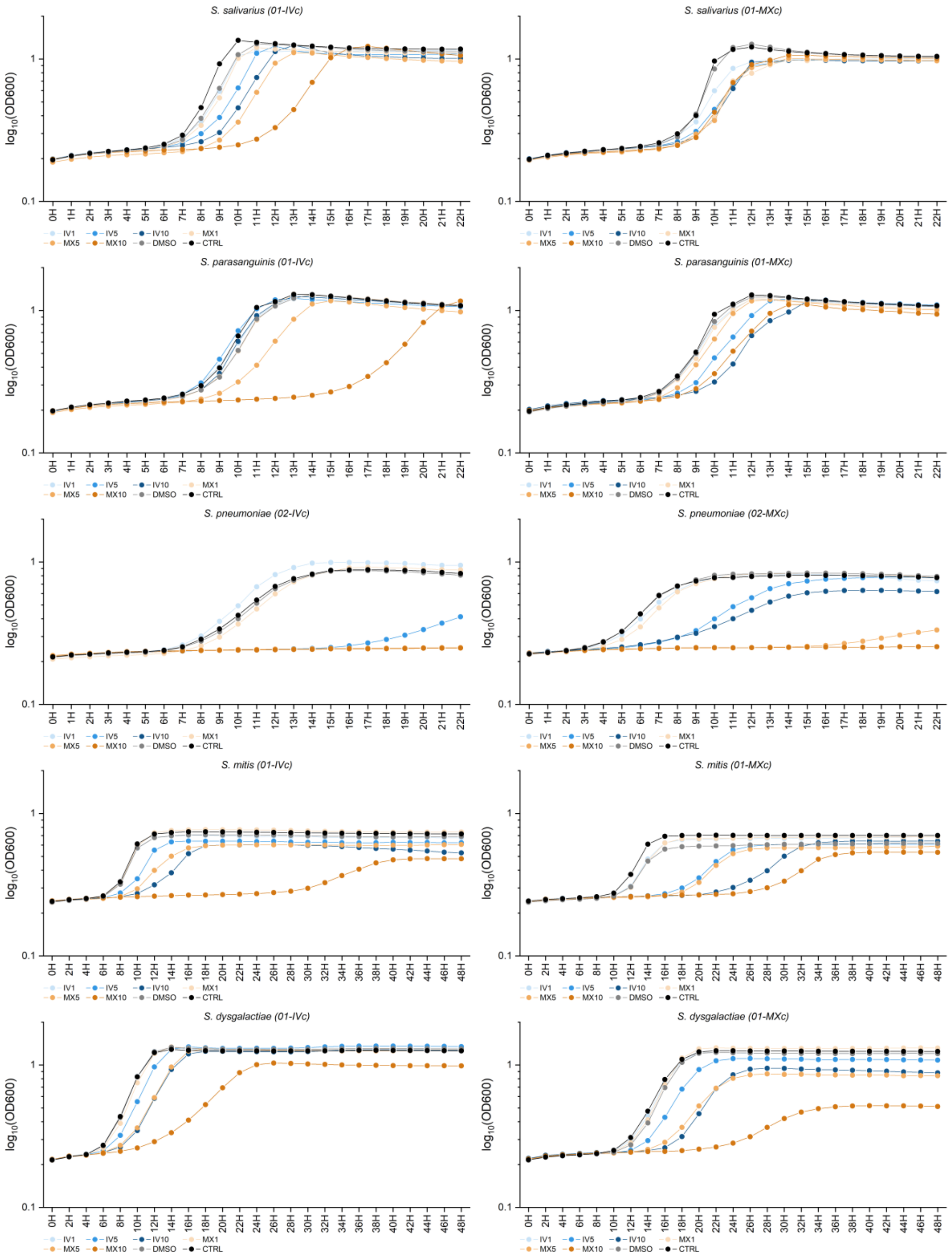


*Supplementary Figure 133: Aerobic growth curves of 10 anthelminthic-prechallenged bacterial isolates co-incubated with IV (blue) or MX (orange) at either 1µM, 5µM or 10µM. Each datapoint represents an average of two independent experiments. The grey curves represent growth in presence of 0.2% DMSO, the black curve represents a positive growth control in presence of BHI + 5% yeast only. Left to right and top to bottom: S. salivarius (01-IVc), S. salivarius (01-MXc), S. parasanguinis (01-IVc), S. parasanguinis (01-MXc), S. pneumoniae (02-IVc), S. pneumoniae (02-MXc), S. mitis (01-IVc), S. mitis (01-MXc), S. dysgalactiae (01-IVc) and S. dysgalactiae (01-MXc).*

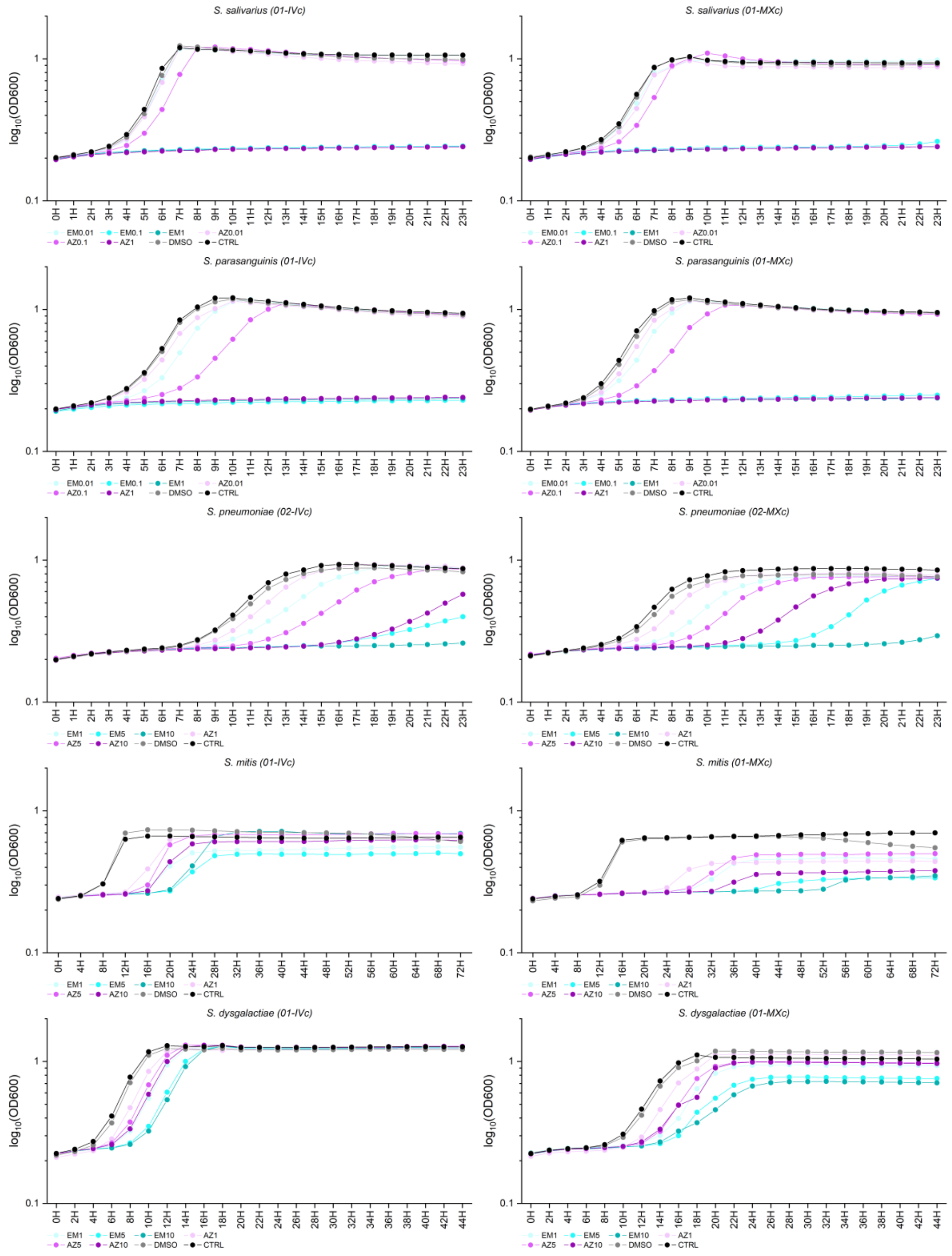


*Supplementary Figure 144: Aerobic growth curves of 10 anthelminthic-prechallenged bacterial isolates co-incubated with EM (turquoise) or AZ (purple) at three different concentrations. Each datapoint represents an average of two independent experiments. The grey curves represent growth in presence of 0.2% DMSO, the black curve represents a positive growth control in presence of BHI + 5% yeast only. Left to right and top to bottom: S. salivarius (01-IVc), S. salivarius (01-MXc), S. parasanguinis (01-IVc), S. parasanuinis (01-MXc), S. pneumoniae (02-IVc), S. pneumoniae (02-MXc), S. mitis (01-IVc), S. mitis (01-MXc), S. dysgalactiae (01-IVc) and S. dysgalactiae (01-MXc).*

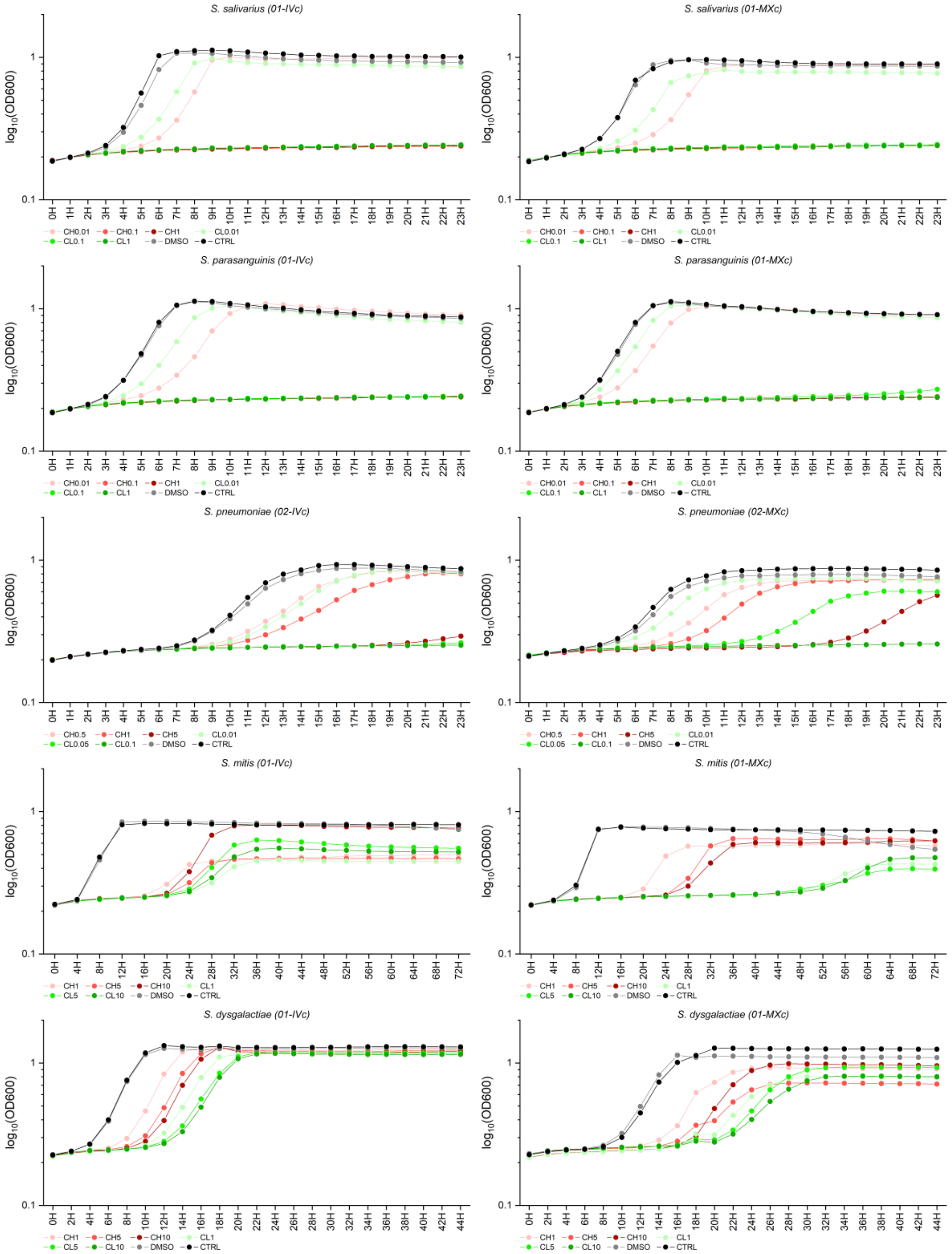


*Supplementary Figure 15: Aerobic growth curves of 10 anthelminthic-prechallenged bacterial isolates co-incubated with CH (red) or CL (green) at three different concentrations. Each datapoint represents an average of two independent experiments. The grey curves represent growth in presence of 0.2% DMSO, the black curve represents a positive growth control in presence of BHI + 5% yeast only. Left to right and top to bottom: S. salivarius (01-IVc), S. salivarius (01-MXc), S. parasanguinis (01-IVc), S. parasanuinis (01-MXc), S. pneumoniae (02-IVc), S. pneumoniae (02-MXc), S. mitis (01-IVc), S. mitis (01-MXc), S. dysgalactiae (01-IVc) and S. dysgalactiae (01-MXc).*

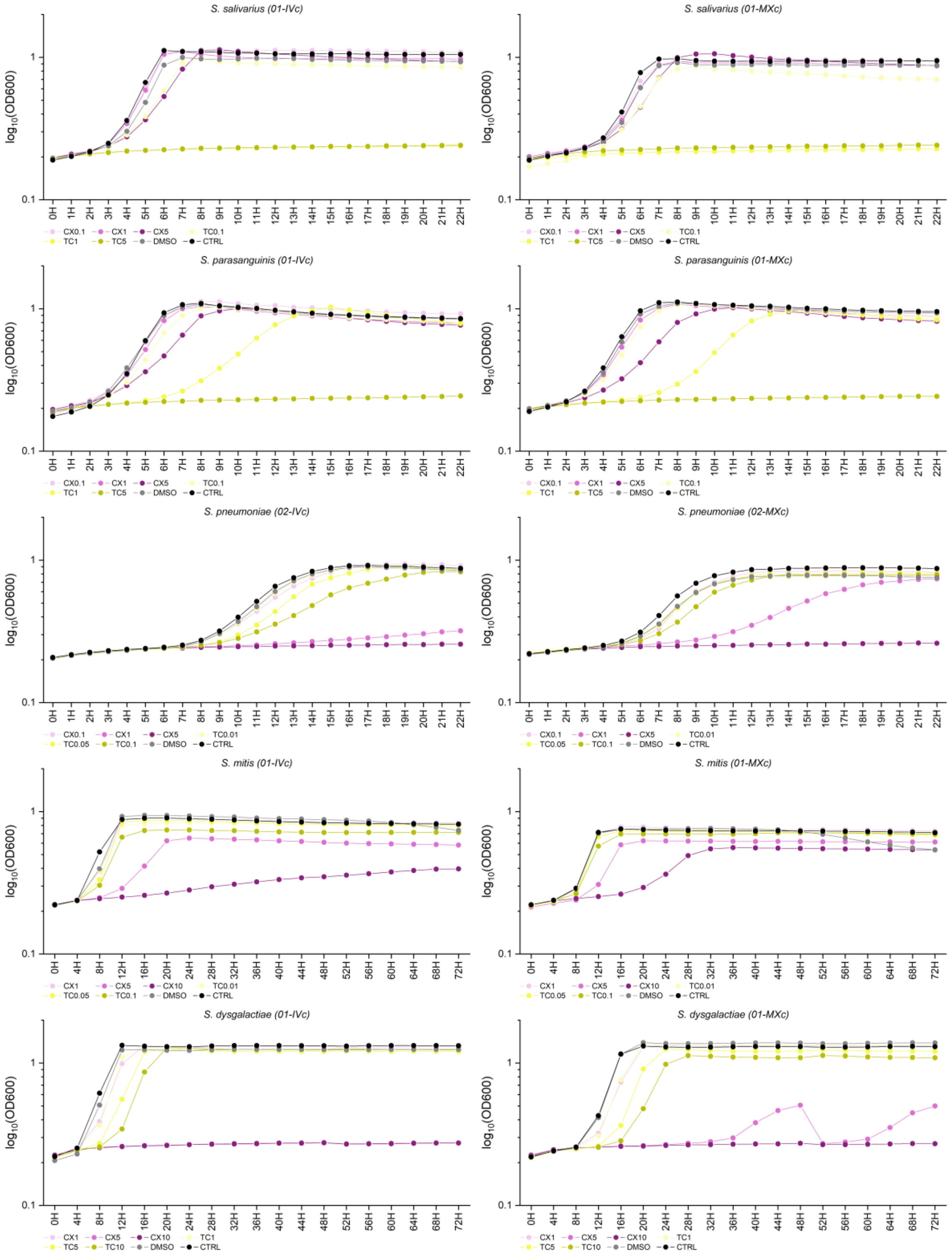


*Supplementary Figure 156: Aerobic growth curves of 10 anthelminthic-prechallenged bacterial isolates co-incubated with CX (mauve) or TC (yellow) at three different concentrations. Each datapoint represents an average of two independent experiments. The grey curves represent growth in presence of 0.2% DMSO, the black curve represents a positive growth control in presence of BHI + 5% yeast only. Left to right and top to bottom: S. salivarius (01-IVc), S. salivarius (01-MXc), S. parasanguinis (01-IVc), S. parasanuinis (01-MXc), S. pneumoniae (02-IVc), S. pneumoniae (02-MXc), S. mitis (01-IVc), S. mitis (01-MXc), S. dysgalactiae (01-IVc) and S. dysgalactiae (01-MXc).*

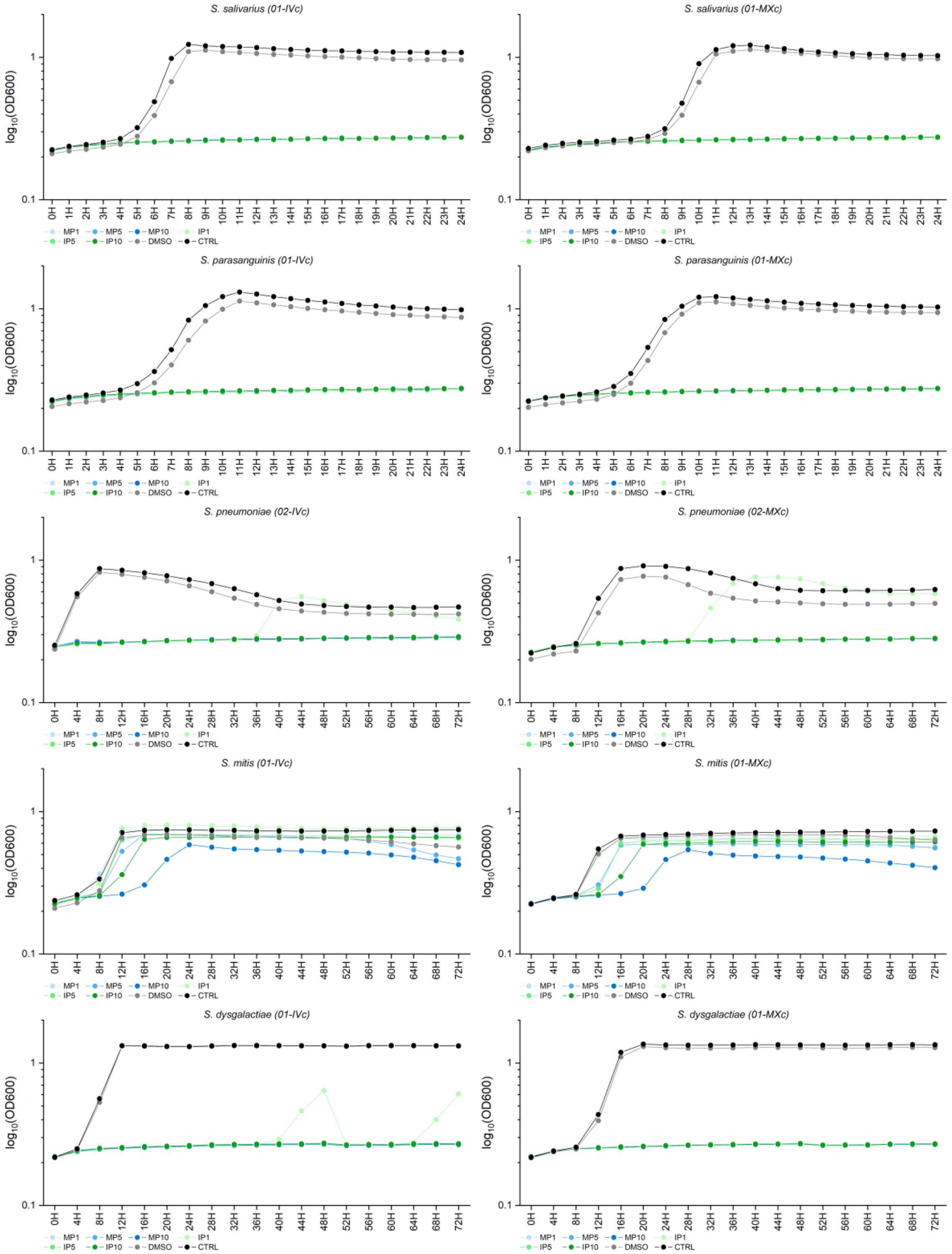


*Supplementary Figure 167: Aerobic growth curves of 10 anthelminthic-prechallenged bacterial isolates co-incubated with MP (blue) or IP (green) at three different concentrations. Each datapoint represents an average of two independent experiments. The grey curves represent growth in presence of 0.2% DMSO, the black curve represents a positive growth control in presence of BHI + 5% yeast only. Left to right and top to bottom: S. salivarius (01-IVc), S. salivarius (01-MXc), S. parasanguinis (01-IVc), S. parasanuinis (01-MXc), S. pneumoniae (02-IVc), S. pneumoniae (02-MXc), S. mitis (01-IVc), S. mitis (01-MXc), S. dysgalactiae (01-IVc) and S. dysgalactiae (01-MXc).*
